# Cerebellar contribution to preparatory activity in motor neocortex

**DOI:** 10.1101/335703

**Authors:** Francois P. Chabrol, Antonin Blot, Thomas D. Mrsic-Flogel

## Abstract

In motor neocortex, preparatory activity predictive of specific movements is maintained by a positive feedback loop with the thalamus. Motor thalamus receives excitatory input from the cerebellum, which learns to generate predictive signals for motor control. The contribution of this pathway to neocortical preparatory signals remains poorly understood. Here we show that in a virtual reality conditioning task, cerebellar output neurons in the dentate nucleus exhibit preparatory activity similar to that in anterolateral motor cortex prior to reward acquisition. Silencing activity in dentate nucleus by photoactivating inhibitory Purkinje cells in the cerebellar cortex caused robust, short-latency suppression of preparatory activity in anterolateral motor cortex. Our results suggest that preparatory activity is controlled by a learned decrease of Purkinje cell firing in advance of reward under supervision of climbing fibre inputs signalling reward delivery. Thus, cerebellar computations exert a powerful influence on preparatory activity in motor neocortex.

## Introduction

Persistent firing, a hallmark of cortical activity in frontal areas of the neocortex (Funahashi et al., 1989; Fuster and Alexander, 1971; Li et al., 2015; Tanji and Evarts, 1976; Wise, 1985), links past events to future actions. In the motor-related areas of the neocortex, the persistent activity that emerges prior to movement is often referred to as preparatory activity (Churchland et al., 2006; Kubota and Hamada, 1979; Paz et al., 2003; Wise, 1985), but the circuit mechanisms underlying the origin, timing and control of this activity remain unclear. A positive thalamic feedback loop has been shown to be involved in the maintenance of preparatory signals in anterolateral motor neocortex (ALM) of mice (Guo et al., 2017), raising the possibility that extra-cortical inputs might regulate neocortical activity in advance of goal-directed movements (Kopec et al., 2015; Ohmae et al., 2017; Tanaka, 2007) via the thalamus. A prominent extra-cortical input to the motor thalamus is provided by the cerebellum (Guo et al., 2017; Ichinohe et al., 2000; Middleton and Strick, 1997; Thach and Jones, 1979), a key brain structure for the learning of sensorimotor and internal context relevant for movement timing (Mauk and Buonomano, 2004). The cerebellum is therefore a plausible candidate for participating in the computation of preparatory activity.

The cerebellum is bidirectionally connected with the neocortex via the disynaptic cerebello-thalamo-cortical and cortico-ponto-cerebellar pathways (Jörntell and Ekerot, 1999; Leergaard et al., 2006; Lu et al., 2007; Proville et al., 2014; Suzuki et al., 2012). The sole output of the cerebellum are the deep cerebellar nuclei (DCN), where ~40 inhibitory Purkinje cells converge on individual postsynaptic neurons (Person and Raman, 2012). The dentate (DN), interpositus (IPN) and fastigial (FN) subdivisions of the deep cerebellar nuclei send excitatory projections to the motor thalamic regions linked to cortical areas involved in the preparation and execution of voluntary movements (Angaut and Bowsher, 1970; Gao et al., 2018; Hoover and Vertes, 2007; Ichinohe et al., 2000; Kelly and Strick, 2003; McCrea et al., 1978; Middleton and Strick, 1997; Sawyer et al., 1994; Shinoda et al., 1985; Thach and Jones, 1979). Although the cerebellum is mostly known for its role in rapid adjustments in the timing and degree of muscle activation, neurons at different stages of cerebellar hierarchy can also represent signals related to upcoming movements or salient events such as reward (Giovannucci et al., 2017; Huang et al., 2013; Kennedy et al., 2014; Wagner et al., 2017). For instance, DN neurons exhibit ramping activity predictive of the timing and direction of the self-initiated saccades (Ashmore and Sommer, 2013; Ohmae et al., 2017). Moreover, inactivation of IPN activity reduces persistent activity in a region of medial prefrontal cortex involved in trace eyeblink conditioning (Siegel and Mauk, 2013). Finally, a recent study has established the existence of a loop between ALM and the cerebellum necessary for the maintenance of preparatory activity (Gao et al., 2018). These results suggest that the cerebellum participates in programming future actions, but the details of how it may contribute to preparatory activity in the neocortex during goal-directed behaviour remain to be determined.

## RESULTS

### Preparatory activity in ALM prior to reward acquisition in a virtual corridor

We developed a visuomotor task in which mice ran through a virtual corridor comprising salient visual cues to reach a defined location where a reward was delivered (80 cm from the appearance of the second checkerboard pattern; Figure 1A; see Methods). Within a week of training, mice learned to estimate the reward location from visual cues, and adjusted their behaviour accordingly, by running speedily through the corridor before decelerating abruptly and often licking in anticipation of reward delivery (Figure 1B,C and Figure S1A-C). This behavioural progress was apparent during the recording session, as the number of false alarm licks outside of the reward zone decreased within tens of trials (Figure S1B), while deceleration and lick onsets emerged in anticipation of reward (Figure S1C).

**Figure 1.**
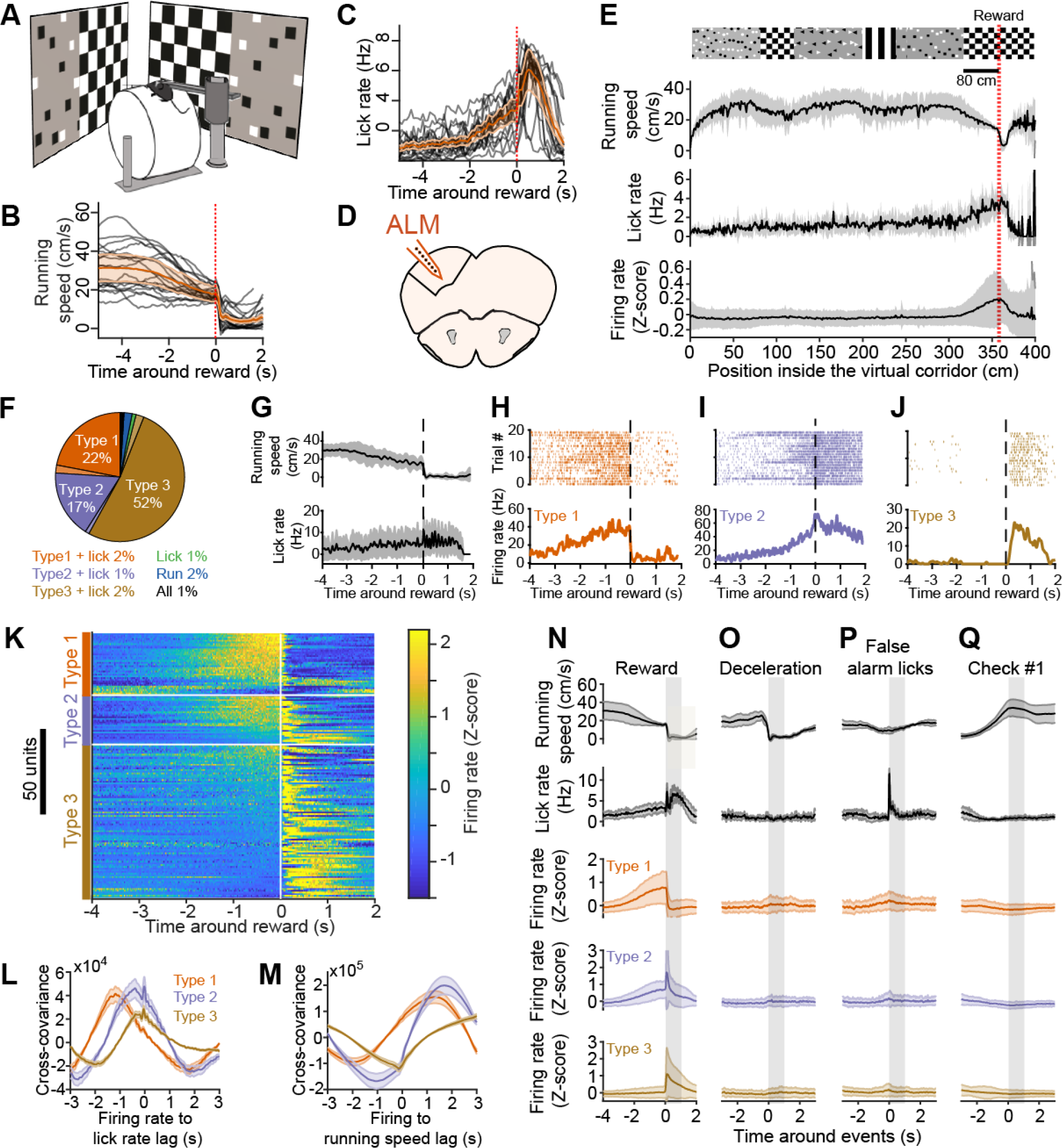
Preparatory activity in the anterolateral motor cortex. Schematic of the virtual reality setup. (B) Running speed profiles for all mice from Figures 1 and 2 (black curves, 21 expert mice) and population average (orange trace, shading is SD). Red vertical dashed line indicates reward. (C) Same as (B) but for lick rate. (D) Schematic showing recording location in the anterolateral motor cortex (ALM). (E) From top to bottom: structure of visual textures lining the virtual corridor walls with the red dotted line indicating the position of reward delivery 80 cm from the appearance of the checkerboard pattern (scale bar), average (black line) and SD (shaded area) of running speed, lick rate and Z-scored firing rate of all ALM neurons exhibiting task-modulation, as a function of position in the virtual corridor. The visual patterns are aligned to the position at which they fully appear in the field of view of the mice (i.e. when they reach the back edge of the monitors). (F) Summary ALM neuron classification (*n* = 169 neurons, 6 mice). (G) Running speed (top) and lick rate (bottom) around reward time for an example recording. Black line is the average, grey shaded area is the SD. (H-J) Spiking activity from example neurons in ALM, classified as type 1, type 2 and type 3 from the same recording as in (G). Top, spike raster for 20 consecutive trials. Bottom, average response profile centered on reward delivery from the same trials shown above. The vertical dotted lines across (G-J) indicate reward time. (K) Mean Z-scored firing rate of reward time-modulated ALM neurons centered on reward time represented by the white vertical line. Within types, neurons (one per line) are sorted by their mean Z score value in the last second before reward. (L) Average cross-covariance between firing rates of all neurons (grouped by type) and lick rate for −2 to 2 s time lags (10 ms binning). The shaded areas represent SEM. (M) Cross-covariance between firing rates and running speed, description as in panel (L). (N) Average (line) and SD (shaded area) centered on reward delivery (vertical shaded area represents time window from reward to reward + 1 s) for, from top to bottom, running speed, lick rate, and firing rate (z-scored), averaged for all type 1, type 2, and type 3 neurons. (O-Q), Same as in (N) for responses centered on deceleration events outside of the reward zone, first lick of a train outside of the reward zone and the appearance of the first non-rewarded checkerboard visual stimulus at the front edge of the monitors (see Figure S3).

We used silicon probes to record the spiking of neurons in ALM neocortex (Figure 1D) to determine how their activity was modulated during our behavioural task, especially during the transition period between running and licking around reward delivery. We found that the activity of task-modulated ALM neurons (*n* = 169, 6 mice; see below) remained, on average, at baseline levels during the entire trial except in the rewarded section of the corridor where it substantially increased (Figure 1E). We identified the neural correlates of running, licking, or reward context by applying a generalised linear model (GLM) (Park et al., 2014) to classify ALM neurons according to running speed, lick times and reward times (Figure S2; see Methods). The activity of 49% of putative ALM pyramidal neurons (see Methods) was modulated by these task variables (*n* = 169/343, 6 mice). The activity of 91% of those units was related to reward times, while that of the remaining units was modulated by running, licks or a combination of these behavioural variables and reward times (Figure 1F). All neuron classes included a minority of units exhibiting decrease rather than increase in activity. The population modulated by reward times included neurons with activity starting a few seconds before reward delivery (referred to as ‘preparatory activity’) and terminating abruptly thereafter (classified as ‘type 1’; see (Svoboda and Li, 2018); *n* = 37, 33 with increasing and 4 with decreasing activity; Figure 1G,H,K), neurons active before and after reward delivery (‘type 2’; *n* = 29, 20 with activity increasing before and after reward, 7 with activity decreasing before and increasing after reward and 2 with activity increasing before and decreasing after reward; Figure 1I,K), or neurons active after reward delivery (‘type 3’; *n* = 88, 79 with increasing and 9 with decreasing activity; Figure 1J,K), consistent with ALM activity observed during a delayed licking task in mice (Svoboda and Li, 2018). Accordingly, ALM population activity tiled the period around reward acquisition (Figure 1K). Cross-covariance analysis between firing rate and lick rate revealed that preparatory activity arises long before the time of the reward (Figure 1L). Type 1-3 neuronal activity in ALM preceded lick rate by 1188, 383, and 350 ms (peaks of mean cross-covariances), respectively, on average. Moreover, type 1 and 2 neuronal activity preceded running speed changes (Figure 1M), albeit with anti-correlation, by 1961 ms and 1045 ms on average, respectively.

To verify that the activity of type 1-3 ALM neurons was specific to reward context (Figure 1N), we examined whether their firing was related to changes in motor output or visual input outside of the rewarded corridor location. Specifically, their activity was not modulated by deceleration events (Figure 1O) or licking bouts outside of the reward zone (Figure 1P), nor by the appearance of non-rewarded checkerboards in a different segment of the virtual corridor (Figure 1Q). These results confirm that the activity of type 1 and 2 ALM neurons before reward acquisition does not reflect motor action or sensory input per se but instead is consistent with preparatory activity building up in anticipation of licking for reward, as described previously (Guo et al., 2014; Li et al., 2015). We noticed that mice consistently decelerated after the appearance of the non-rewarded checkerboard (Figure 1E and Figure S3A), although they did not produce substantially more licks on average at this corridor position (Figure 1E and Figure S3B). To verify whether we could see any correlate of this behaviour in ALM activity (Figure S3C), we plotted the activity of type 1-3 neurons as a function of distance in the corridor (Figure S3D). Type 1-3 neurons were selectively active around the position of reward delivery, showing very little modulation of activity around the non-rewarded checkerboard (Figure S3D,E).

A likely role for preparatory activity is to anticipate the timing of future events based on past experience and environmental cues in order to produce accurately timed actions (Mauritz and Wise, 1986; Roux et al., 2003; Tsujimoto and Sawaguchi, 2005). If ALM preparatory activity is related to predicting the timing of future rewards, the 2 following conditions should be met. First, the slope of preparatory activity should depend on the delay between the environmental cues and the reward, that is if this delay is long, preparatory activity should start early and progressively increase until reward delivery and if the delay is short preparatory activity should exhibit a steeper ramp. Second, the onset of preparatory activity should start at the appearance of the environmental cue regardless of the delay to reward delivery. As in our task the delay between environmental cues and rewards depends on the mouse speed, we grouped trials according to mouse deceleration onset before rewards (Figure S4). The onset of increase in activity in type 1 and 2 neuronal populations was closely related to the deceleration profiles, starting earlier in trials when mice decelerated sooner in anticipation of reward (Figure S4A-C). Accordingly, in trials where the mice decelerated closer to rewards, preparatory activity started later and exhibited a steeper ramp. In our task, the environmental cue signifying the future occurrence of rewards is likely to be the appearance of the final checkerboard (Figure 1E). When plotting the same groups of trials around the appearance of the rewarded checkerboard, the onsets of deceleration and increase in type 1 and 2 neuronal activity largely overlapped, starting around 1 second after the appearance of the rewarded checkboard when mice have fully entered the reward zone (Figure S4D,E). Taken together these data strongly support the view that ALM preparatory activity tracks the passage of time from environmental cues in order to accurately time motor preparation to obtain the reward.

### The cerebellar dentate nucleus exhibits preparatory activity

Since the DN sends excitatory projections to the motor thalamus (Guo et al., 2017; Ichinohe et al., 2000; Middleton and Strick, 1997; Thach and Jones, 1979), which has been shown to participate in the maintenance of preparatory activity in mouse ALM neocortex (Guo et al., 2017), we investigated whether DN activity could influence ALM processing. We first recorded the activity of DN neurons to determine how their activity was modulated during the task (Figure 2A). Forty four percent of all recorded DN neurons (*n* = 355, 15 mice) could be classified according to our task variables and the activity of 69% of classified DN neurons was related to reward times only (Figure 2B). The activity of the other neurons was related to lick times, running, or a mixture of these variables plus reward times (Figure 2B). Of neurons whose activity was modulated by reward times only, 13% were classified as type 1 (*n* = 20, 18 with increasing and 2 with decreasing activity), 20% as type 2 (*n* = 32, 22 with activity increasing before and after reward, 9 with activity decreasing before and increasing after reward and 1 with activity increasing before and decreasing after reward), and 36% as type 3 neurons (*n* = 57, 49 with increasing and 6 with decreasing activity; Figure 2B-G). As in ALM, type 1-3 neuronal activity tiled the period around reward delivery (Figure 2G). Type 1, 2, and 3 neurons’ spiking preceded lick rate by 915, 50 and 10 ms (peaks of mean cross-covariances), respectively, on average (Figure 2H). Moreover, type 1 DN neuron activity was anti-correlated with running speed and preceded its changes by 1972 ms on average (Figure 2I). Type 2 and 3 DN neuronal activity emerged less in advance of behavioural changes on average compared to ALM (Figure 1L,M), although the distribution of cross-covariance peak times were not significantly different between ALM and DN neurons (*p* > 0.05, Wilcoxon signed-rank test). As in ALM, the activity of type 1-3 DN neurons was not modulated by changes in motor behaviour or visual input (Figure 2K-M), but specifically emerged around the position of reward delivery (Figure 2J and Figure S3F-H). Also akin to ALM activity was the dependency of the duration of type 1 DN neuronal activity on the delay between mice decelerations and reward times, which was less apparent for the population of type 2 DN neurons (Figure S4F-H). As for ALM preparatory activity, the onset of type 1 DN neuronal activity was well timed once mice entered the rewarded part of the corridor (Figure S4I,J), supporting the view that preparatory activity keeps track of time between visual cues and rewards. Thus, preparatory activity in DN (type 1 neurons) was largely indistinguishable from that recorded in ALM. Type 2 and 3 DN neuronal populations on the other hand seemed to be more closely related to behavioural changes than their counterparts in ALM (see cross-covariance plots in Figure 2H,I vs Figure 1L,M and Figure S4). To test whether preparatory activity is somewhat specific to DN or can also be found in other deep cerebellar nuclei, we recorded from the IPN (Figure S5C), located more medially than the DN (Figure S5A), which also projects to the motor thalamus (Guo et al., 2017). We found a lower proportion of neurons classified as type 1 (1/74 classified neurons, 5 mice) and type 2 (8/74) in the IPN compared to DN, although this difference was only significant for type 1 neurons (Figure S5B,D-F). On the other hand, type 3 neurons were more enriched in the IPN (54/74 classified neurons) than in the DN population (Figure S5B,D-F). Hence, preparatory activity appears to be more represented in the DN.

**Figure 2.**
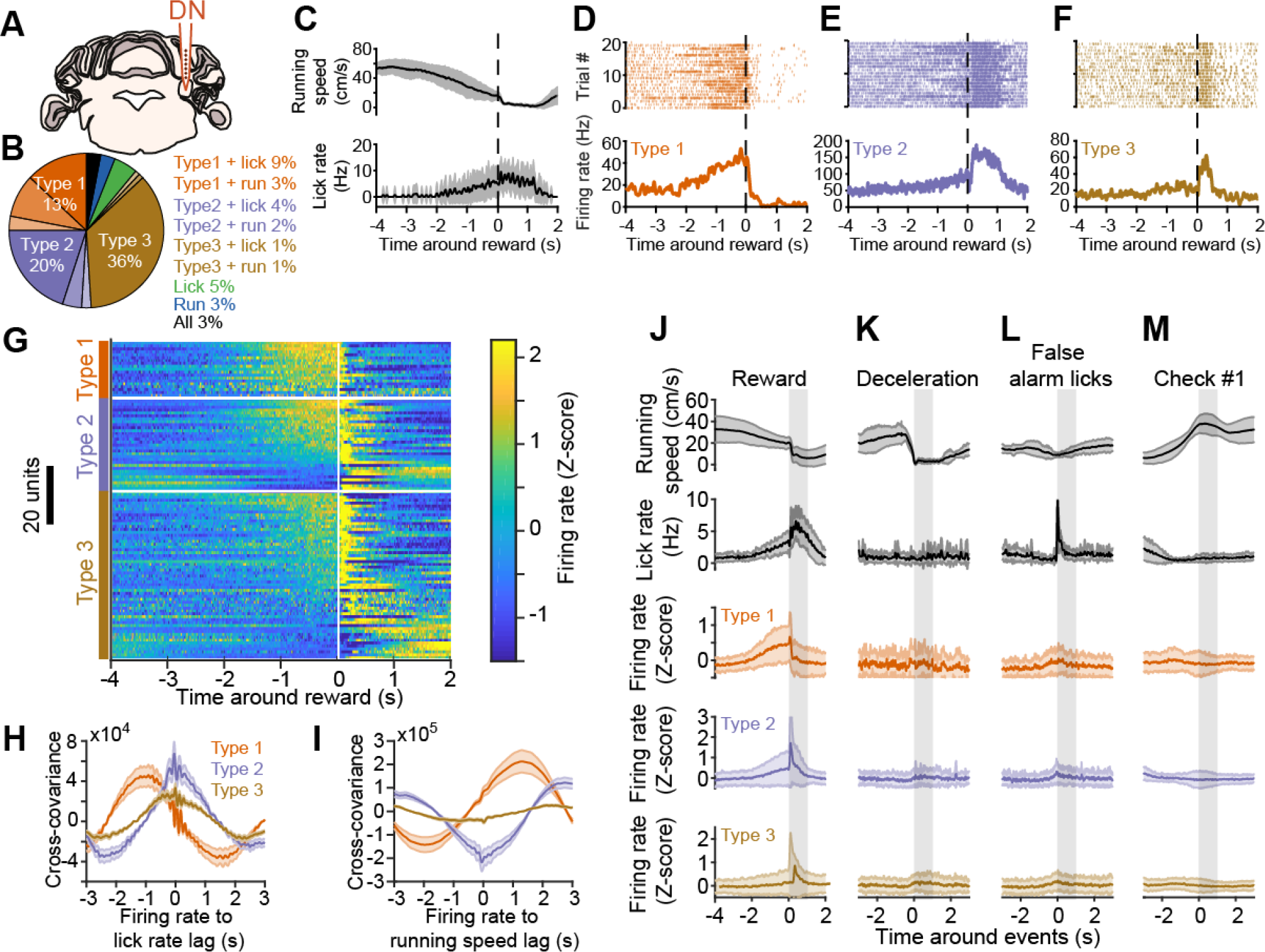
Preparatory activity in the dentate nucleus. (A) Schematic showing recording location in the dentate nucleus (DN). (B) Summary DN neuron classification (*n* = 156 neurons, 15 mice). (C) Running speed (top) and lick rate (bottom) around reward time for an example recording. Black line is the average, grey shaded area is the SD. (D-F) Spiking activity from example neurons in DN, classified as type 1, type 2 and type 3 (same recording as in (C)). Top, spike raster for 20 consecutive trials. Bottom, average response profile centered on reward delivery from the same trials shown above. The vertical dotted lines across (C-F) indicate reward time. (G) Activity of reward-modulated DN neurons centered on reward time (white vertical line). Within types, neurons (one per line) are sorted by their mean Z score value in the last second before reward. (H) Average cross-covariance between firing rates of all neurons (grouped by type) and lick rate for −2 to 2 s time lags (10 ms binning). The shaded areas represent SEM. (I) Cross-covariance between firing rates and running speed, description as in panel (H). (J) Average (line) and standard deviation (shaded area) centered on reward delivery (vertical shaded area represents time window from reward to reward + 1 s) for, from top to bottom, running speed, lick rate, and firing rate (z-scored), averaged for all type 1, type 2, and type 3 units. (K-M), Same as in (J) for responses centered on deceleration events outside of the reward zone, first lick of a train outside of the reward zone and the appearance of the first non-rewarded checkerboard visual stimulus at the front edge of the monitors (see Figure S3).

### DN participates to ALM preparatory activity

To determine the contribution of DN firing on ALM preparatory activity, we silenced DN output by photoactivating cerebellar Purkinje cells (PCs) expressing channelrhodopsin-2 under the control of the specific Purkinje cell protein (PCP2) promoter (Figure 3A; see Methods). Since in our behavioural task reward delivery is dependent on visuomotor context, we targeted the photoactivation to the lateral part of crus 1 in the cerebellar cortex, a region that projects to DN (Payne, 1983; Voogd, 2014) and uniquely responds to electrical stimulation of both visual and motor cortex (Figure S6), suggesting it integrates visuomotor signals from the neocortex. The activation of PCs in lateral crus 1 began 20 cm in advance of the rewarded position in the virtual corridor (Figure 1E) and lasted 1 s in order to terminate around reward delivery. Simultaneous silicone probe recordings from DN and ALM (Figure 3A) revealed that optogenetic activation of PCs effectively silenced most DN neurons (Figure 3B and Figure S7A) regardless of response type (Figure 3L,M; firing rate control vs PC photoactivation: 49.7 +/− 33.4 Hz vs 4.1 +/− 10 Hz, 92 % decrease, *p* < 0.0001, *n* = 69, 3 mice), consequently resulting in a substantial reduction of activity in a large fraction of ALM neurons (*n* = 98/279, 5 mice; Figure 3C-K and Figure S7B). Activity of all type 1 and most type 2 ALM neurons was robustly decreased by PC photoactivation (respectively, 14/14 and 34/49 neurons, Figure 3N-P), such that, on average, their firing rate decreased to baseline activity levels (type 1 control: 18.5 +/− 14 Hz vs photoactivation: 9.8 +/− 12.7 Hz, *n* = 14, *p* = 0.0001; type 2 control: 20.3 +/− 16.1 Hz vs photoactivation: 11.5 +/− 11.7 Hz, *n* = 49, *p* < 0.0001, Figure 3H,I,N,O). Type 3 and unclassified ALM neurons exhibited a mixture of effects, including a fraction of units that were excited (Figure 3E,P), and their population activity during PC photoactivation was not affected on average (respectively 10.8 +/− 8.9 Hz vs 9.9 +/− 9.3 Hz, *p* = 0.09, *n* = 74 and 9.8 +/− 9.8 Hz vs 10.2 +/− 10.3 Hz, *p* = 0.89, *n* = 165; Figure 3J,K,N,O). Type 1 and 2 neurons were significantly more inhibited during photoactivation (Figure S8A) and had higher firing rates (Figure S8B) than type 3 and unclassified neurons. The effect of photoactivation did not seem to be explained by the difference of firing rate between neuron types (Figure S8C-F) and, even when excluding neurons with low firing rate (below 10 Hz), only type 1 and type 2 neurons were significantly affected by photoactivation (Figure S8G). Furthermore, the amplitude of the effect of PC photoactivation appeared to be dependent on the size of the ramp (as defined in Figure S8H) and not on the firing rate in control condition (Figure S8I). Indeed, in a linear model including ramp size and control firing rate (see Methods), the photoactivation effect was proportional to the ramp size and not on the firing rate (*p* < 1e-5 vs *p* = 0.7). Therefore, lateral crus 1 PC photoactivation preferentially impacted ALM neurons whose activity was modulated in anticipation of reward.

**Figure 3.**
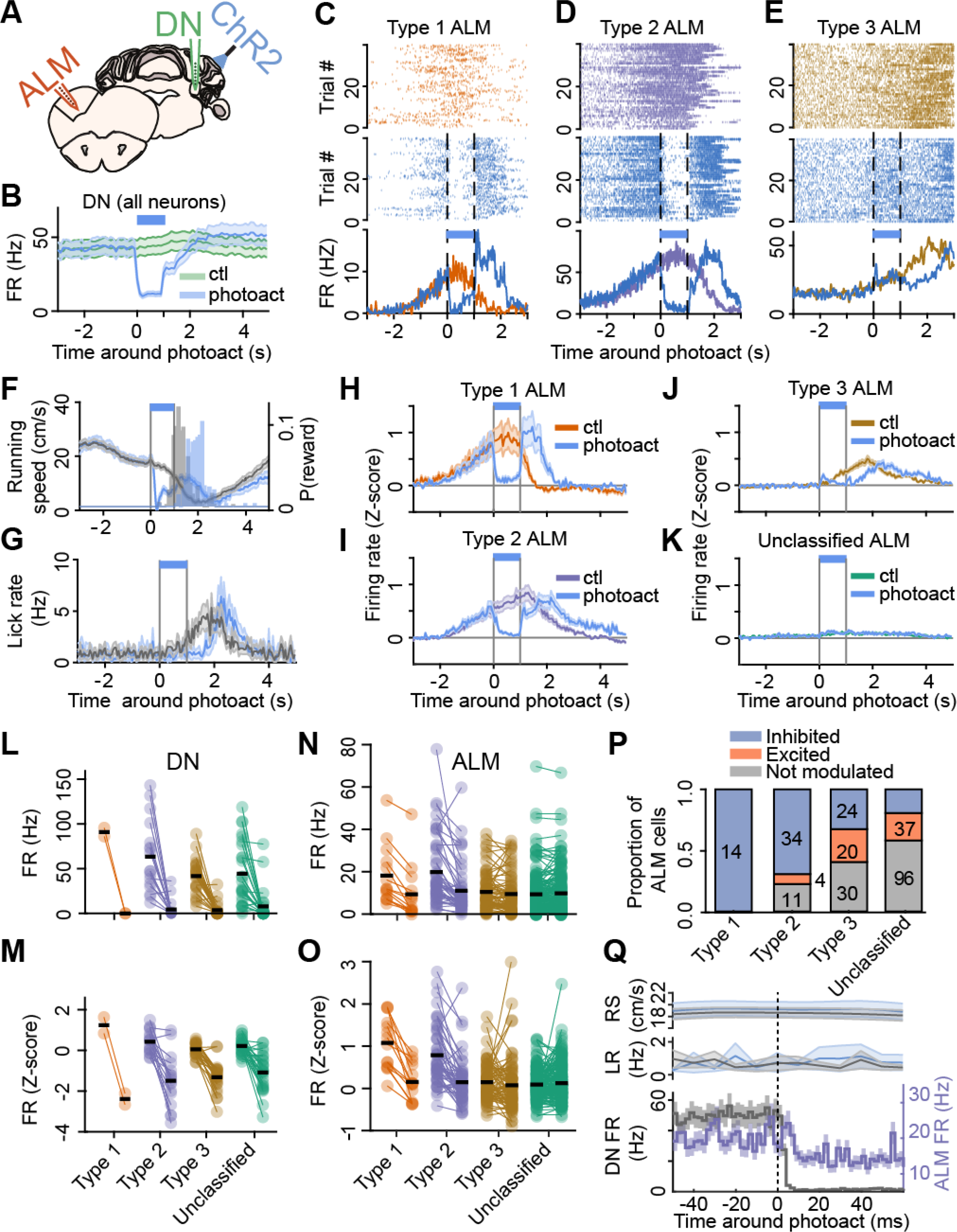
Cerebellar output is required for the persistence of ALM preparatory activity. Schematic of experiments. The dentate nucleus (DN, green) and anterolateral motor cortex (ALM, orange) were simultaneously recorded in L7-ChR2 mice performing the task during photoactivation of Purkinje cells in lateral crus 1 (PCs, blue light). (B) PC photoactivation effectively silenced DN population activity. Duration of photoactivation is indicated by a blue bar. (C-E) Raster plot during control trials (top) and photoactivation trials (middle) and average responses (bottom) from example neurons recorded in ALM classified as type 1 (C), type 2 (D) or type 3 (E). (F-G) Average profiles around time of stimulation in photoactivation trials (blue traces, average and SD) and control trials (grey) for running speed (F) and lick rate (G) and distribution of reward probability (P(reward)) in photoactivation trials (F, blue histogram) and control trials (F, grey histogram). (H-K), Response profiles of ALM neurons aligned to photoactivation onset for type 1 (H), type 2 (I), type 3 (J) and unclassified cells (K) during photoactivation (blue traces, average and SD) and control trials (coloured traces). (L-O) Quantification of modulation. Average firing rate (L,N) and z-scored firing rate (M,O) during the first second after stimulation time for DN neurons (L,M) and ALM neurons (N,O). For all plots the first column of dots (one per cell) is the control condition, the second column the photoactivation condition. The black lines indicate the population mean. All types of DN cells were inhibited by photoactivation (L,M) while only type 1 and 2 ALM cells showed a significant inhibition (N,O). (P) Proportion of cells being inhibited (blue), excited (orange) or not significantly modulated (grey) for the 4 classes of ALM neurons (number of cells indicated in each bar, except for excited type 2 neurons where it is shown on the right side of the bar). (Q) Average response profiles of firing rate for all DN neurons (grey) and all type 1 and type 2 ALM neurons (purple) with significant reduction in activity compared to control trials, during the first 60 ms following PC photostimulation onset (vertical dotted line). Traces with shaded areas are mean +/− SEM. Running speed (RS) and lick rate (LR) did not change in this time window and were not significantly different between control (black, average and SD) and photoactivation (blue) trials. Data shown in panels (L-Q) include an additional recording (6 recordings in total, 5 mice) where the photoactivation period lasted 2 seconds (versus 1 second in all other recordings).

In trials with PC photoactivation, mice transiently ceased to run approximately 170 ms after light onset (Figure 3F, curve; Figure S7C), reaching the reward later (Figure 3F, histograms) and licking at that time (Figure 3G). This effect on running is reminiscent of dystonic postures that have been previously observed during cerebellar deficits or pharmacological activations (Shakkottai, 2014) and prevents any interpretation of the effect of PCs photoactivation on preparatory behaviour. As type 3 neurons were modulated after the reward, their firing peaked later in photoactivation trials (Figure 3J) but remained aligned to reward delivery (Figure S9). However, the suppression of neuronal activity in DN and ALM was not a consequence of running speed change because the onset of the firing rate decrease was almost immediate in DN neurons (2 ms; see Methods) and was 10 ms for type 1 and 2 ALM neurons (Figure 3Q) and a significant decrease in ALM activity during PC photoactivation was observed before any change in running speed (10-150 ms after photoactivation onset; type 1 control: 18.3 +/− 14.5 Hz vs photoactivation: 10.2 +/− 14.1, *n* = 14, *p* = 0.0017; type 2 control: 19.19 +/− 15.8 Hz vs photoactivation: 14.6 +/− 16.2, *n* = 49, *p* = 0.0002; control running speed: 18.7 +/− 7.5 cm/s vs photoactivation 18.9 +/− 6.3, *p* = 0.84). The short-latency decrease in ALM activity by PC photoactivation (8 ms, accounting for the delay in DN inhibition) was consistent with the time delay expected for the withdrawal of excitation via the disynaptic pathway from DN to ALM via the thalamus. Type 3 ALM neurons that were inhibited exhibited a similar time profile than type 1-2 neurons, with a significant drop in activity 8 milliseconds after PC photoactivation onset (Figure S7D). On the other hand, excited type 3 neurons exhibited significant changes in activity only after 40 milliseconds (Figure S7E), suggesting the involvement of an additional, possibly intracortical, circuit. These results demonstrate that the maintenance of preparatory activity in ALM requires short-latency, excitatory drive from the cerebellum.

Preparatory activity recovered in ALM shortly after the end of PC photoactivation (Figure 3H,I), which suggests the involvement of other brain regions in its maintenance. We tested whether the contralateral cerebellar-cortical circuit, that should remain unaffected during unilateral photoactivation, reinstated ALM activity by photoactivating lateral crus 1 PCs on both sides (Figure S10A). Moreover, we established a progressive ramp in the offset of the laser to avoid activity rebound in the DCN (see Methods). We found no difference in the effect of unilateral vs bilateral PC photoactivation on type 1 (*n* = 10, 8 inhibited, 3 mice), type 2 (*n* = 13, 10 inhibited) or type 3 ALM neurons (*n* = 33, 18 inhibited, 11 excited; Figure S10B), except for shorter latency of inhibition upon unilateral photoactivation (8 ms vs 14 ms for unilateral photoactivation; Figure S10C). These data suggest that other brain regions involved in motor preparation such as the basal ganglia (Kunimatsu et al., 2018), may contribute to the recovery of preparatory activity.

Finally, we tested the role of other deep cerebellar nuclei towards preparatory activity during our task. In contrast to DN, the activity of neurons in IPN was modulated more after reward delivery than before, with fewer neurons exhibiting preparatory activity in anticipation of reward acquisition (Figure S5). Moreover, by photoactivating PCs in lobule IV/V (Figure S11A), which exclusively inhibits neurons in the FN (Voogd, 2014), we observed a strong suppression of type 1 (*n* = 9/10, 3 mice) and type 2 ALM neurons (*n* = 11/15), but with a substantially longer latency than upon lateral crus 1 PC photoactivation (Figure S11B,C). In contrast to the very short latency suppression following lateral crus 1 photoactivation (10 ms; Figure 3), the first significant reduction in type 1-2 ALM during lobule IV/V PC photoactivation occurred after 280 milliseconds (Figure S11D), a delay too long to result from a direct connection from FN to the motor thalamus. Both lateral crus 1 and lobule IV/V PC photoactivation induced a sharp deceleration in mouse running (at 165 and 210 milliseconds respectively), however, unlike for lateral crus 1 PC photoactivation, the behavioural effect during lobule IV/V PC photoactivation preceded the inhibition of type 1-2 ALM neuronal activity, suggesting that the latter resulted from the mouse arrest. Additionally, type 3 ALM neurons appeared more excited upon lobule IV/V PC photoactivation (Figure S11C) but with a similar time profile than with lateral crus 1 PC photoactivation (50 ms onset), indicating that this excitatory effect on ALM activity and the inhibition of running behaviour are not specific to the DN-ALM circuit. Taken together, our results suggest the existence of a dedicated DN output participating in preparatory activity.

### Reward time-based PC learning in lateral crus 1 as a likely mechanism for setting the timing of preparatory activity

To gain insight in how preparatory activity could emerge in DN neurons that are under the inhibitory control of the cerebellar cortex, we recorded simultaneously from putative Purkinje cells (PCs) in lateral crus 1 (Figure S12A-F; see Methods) and from DN neurons (*n* = 3 mice; Figure 4A,B). The firing of PCs was modulated on the same time scale around the time of reward delivery as simultaneously recorded DN neurons (Figure 4C-H). Many PCs exhibited inverse modulation of activity compared to DN neurons, resulting in negative cross-covariances between simultaneously recorded PC-DN pairs (Figure 4I; see Methods), consistent with the fact that PCs provide inhibitory input onto DN neurons. Most PCs ramped down their activity prior to reward (Figure 4K), while DN neurons exhibited either activity increases or decreases (Figure 4J). On average, PCs decreased their firing in the second preceding reward (−0.14 +/− 0.41, *n* = 56, *p* = 0.0005), in contrast to DN neurons (0.06 +/− 0.54, *n* = 69, *p* = 0.4, PC vs DN: *p* = 0.003, Figure 4L). To verify whether this PC activity profile is specific to lateral crus 1 we recorded from the adjacent crus 2 (Figure S13), a cerebellar cortical region which also projects to the DN. We found that crus 2 PCs mostly exhibited decreases in activity after reward (Figure S13E,F), pre-reward decrease in activity being significantly less apparent than in crus 1 (Figure S13G). To test if the recorded regions of crus 1 and DN were functionally connected, we computed the cross-correlogram between all pairs of simultaneously recorded neurons. A small fraction of these correlograms were significantly modulated (46/1855; see Methods; Figure 4M). Six of these modulated pairs showed positive cross-correlations before the spike of the PC (Figure S14A,B). These are likely caused by common inputs exciting both the DN neuron and the PC. The other 40 pairs exhibited a millisecond-latency trough, consistent with monosynaptic inhibition from PC to DN neurons (Figure 4M and Figure S14A-C). Over longer timescale, the average cross-correlogram between all putatively connected pairs (Figure 4N; see Methods) revealed that PC inhibition onto DN neurons is fast, strong, and long-lasting. Thus, the ramping down of PC activity can relieve DN neurons of inhibition and allow extra-cerebellar inputs to drive their ramping activity in anticipation of reward (Figure S14D). Finally, we found that the correlation between the response profiles of the putatively connected PC-DN neuron pairs were not linked to the strengths of their connection (i.e. PCs with positively correlated activity to that of DN neurons also substantially participated to their inhibition; Figure S14E; *p* = 0.92, F-test). Therefore, a learned decrease in the activity of ramping down PCs, rather than plasticity at the PC-DN neurons synapses might explain the emergence of DN preparatory activity.

**Figure 4.**
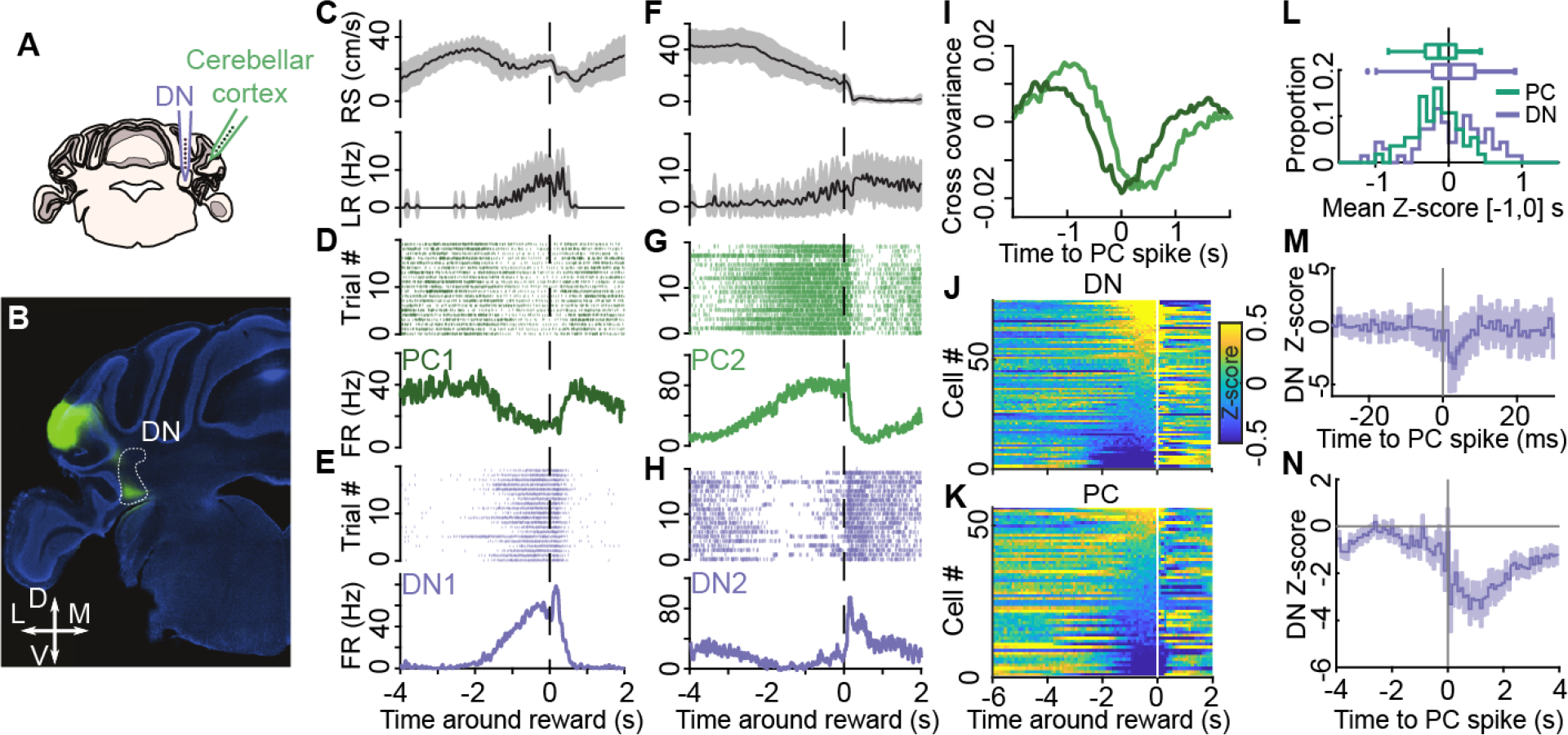
The relationship between activity of lateral crus 1 Purkinje cells and dentate nucleus neurons. Schematic of experiments. The neurons in dentate nucleus (DN, purple) and Purkinje cells (PCs) in the cerebellar cortex (lateral crus 1, green) were simultaneously recorded in mice performing the task. Injection of AAV expressing GFP in the cerebellar cortex marking the axons of PCs (green) projecting to the part of the DN (white outline) that was targeted for recordings. Coronal slice, counterstained with DAPI (blue). (C-H) Examples of two recordings (first recording from (C) to (E), second recording from (F) to (H)) in which PCs and DN neurons were simultaneously recorded. (C,F) Running speed (RS, top) and lick rate (LR, bottom), mean (black line) +/− SD (shaded area). (D,G) spike raster plot (top) and mean firing rate of the same PCs (bottom). (E,H) same as in (D,G) for simultaneously recorded DN neurons. (I) Cross-covariance between the activity of PCs and DN neurons from panel (D,E, dark green) and panel (G,H, light green) at different lags (5ms bins). (J,K) Average response profiles for all DN neurons (J) and all PCs (K) sorted by their mean Z-score value in the last second before reward. White vertical line indicates reward time. (L) Distribution (bottom) and bar plot (top) of mean Z-score value in the last second before reward for PCs (green) and DN neurons (purple). (M) Average Z-scored cross-correlogram for all modulated DN-PC pairs (46/1855 pairs; see Methods) showing a short latency inhibition consistent with a monosynaptic inhibitory connection from PCs to DN neurons. Line is the mean, light purple shading is SD. (N) Average shuffle-corrected cross-correlogram (see Methods) for all putatively connected pairs showing a decrease in the probability of DN neuron firing in the seconds following PC activity. Trace is mean +/− SD.

The best described form of plasticity in the cerebellar cortex is the long-term depression of parallel-fibre to PC synapses under the control of teaching signals conveyed by climbing fibres (Lisberger et al., 1996). To address whether PC activity in lateral crus 1 could be related to such a mechanism (Figure 5A), we first analysed more closely the population of PCs exhibiting a decrease in activity before reward (*n* = 37/72, 4 mice; see Methods). Activity of all PCs was modulated during running bouts and some exhibited tuning to running speed (and/or visual flow speed), following linear or more complex relationships (Figure 5B). By fitting tuning curves of PC activity to running speed we fit linear firing model for each PC (see Methods). The linear model captured well the mean PC firing rates during deceleration events outside of the reward zone (Figure 5C,D,G). However, in comparison, the model significantly overestimated firing around rewards in most PCs (*n* = 25/37; Figure 5E-G; see Methods). This suggests that the strong decrease in the activity of most PCs prior to reward results from learned reductions in firing associated with the reward context. We thus looked for signatures of climbing fibre activity around reward times (Figure 5H). Climbing fibre discharges result in PC complex spikes (Figure S12D-F) that are also apparent as so-called fat spikes likely resulting from the large inward currents occurring in PC dendrites (Gao et al., 2012). We found that fat spikes were readily detectable in our recordings (Figure S12G,H), firing at characteristically low firing frequencies of climbing fibre discharges (Figure S12I) (Zhou et al., 2014). Interestingly, 13/26 of the fat spikes units recorded in lateral crus 1 but 0/26 recorded in crus 2, exhibited a significant increase in their probability of firing (0.61 +/− 0.15; Figure 5I-K; see Methods) shortly after reward delivery (177 +/− 117 ms; Figure 5J,L), consistent with previous reports (Heffley et al., 2018; Ohmae and Medina, 2015). Moreover, the first fat spikes occurring after reward delivery exhibited relatively low jitter (29 +/− 8 ms; Figure 5J,M), which may partly result from the variability in reward consumption across trials. Finally, plotting fat-spike firing rates as a function of position inside the virtual corridor confirmed that they occurred most consistently shortly after rewards (Figure S12J,K). Thus, these putative climbing fibre events are well suited to report the timing of reward delivery to PCs and thereby shape the preceding decrease in their activity.

**Figure 5.**
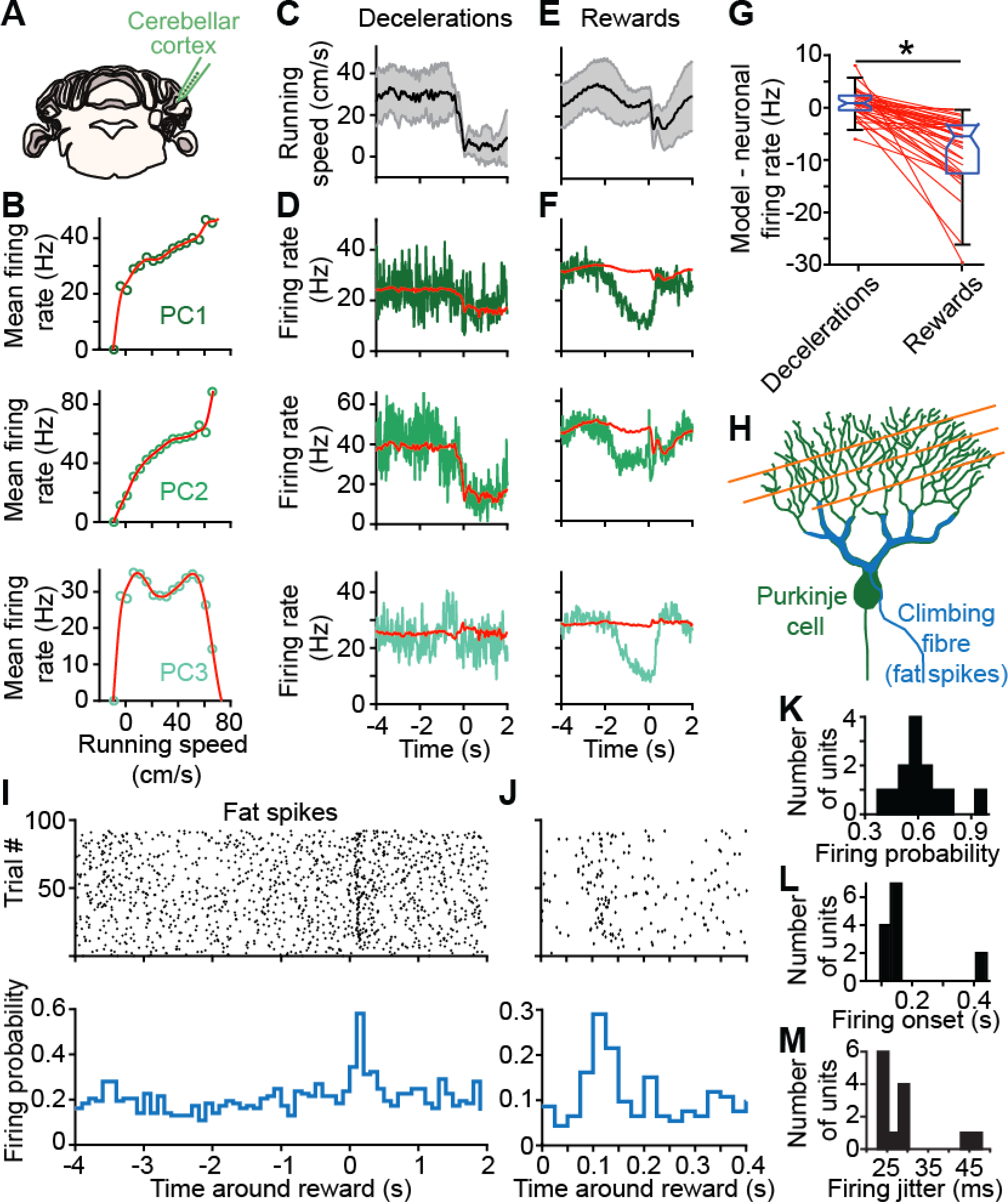
Evidence for reward time-based supervised learning in lateral crus 1 Purkinje cells. (A) Schematic of experiments. Purkinje cell (PC) and climbing fibre-related activity was recorded from lateral crus 1. (B) Plot of firing rates as a function of running speed for 3 simultaneously recorded PCs. Green dots represent average firing rate for a given running speed bin (5 cm/s), the red line is the fit of the tuning curve obtained with a smoothing spline. (C) Mean running speed (black line) and SD (grey shaded area) and (D) mean firing rate of the 3 example PCs (green) and modelled firing rates obtained from the running speed tuning curves (red) shown in (B) around deceleration events outside of the reward zone (t = 0s). (E,F) Running speed and firing profile of the same PCs as in (D) centred around rewards (t = 0s). Note the substantial deviation of firing rates from the model specific to the reward context. (G) Summary plot of mean remaining firing rate after subtracting the modelled activity for all PCs exhibiting decreased activity before reward (see Methods) during the last 2 seconds before the events. Box plots represent quartiles (non-outlier minimum, 25%, median, 75%, and non-outlier maximum values). Paired values from single neurons (red dots) in the decelerations (left column) and rewards (right column) conditions are linked by red lines. PCs activity is significantly lower than that of the model in the reward context (p < 0.0005, Wilcoxon signed-rank test). (H) Schematic of a PC (green) and its two sources of inputs: parallel fibres (orange) which give rise to simple spikes, and the climbing fibre (blue) which give rises to complex spikes also apparent as fat spikes (FS; see Figure S11). (I) Spike raster plot (top) and firing probability (100 ms binning, bottom) aligned on reward (t = 0s) for an example FS unit. (J) Same as (I) but focusing on the time period following reward with 25 ms binning. (K) Distribution of firing probability for all FS units exhibiting significant increase in activity in the 500 ms after reward delivery (*n* = 13, 4 mice; see Methods). (L) Same as in (K) for firing onset (time of first spike in the first 100 ms bin of significant probability increase). (M) Same as in (K) for firing jitter (mean difference between first spike times shown in L).

Finally, we found that the reduction in PC firing occurred specifically in the reward context within the virtual corridor (Figure S15A) and that it exhibited the characteristic cue-to-reward dependent duration and cue-to-reward independent onset properties observed of ALM and DN preparatory activity (Figure S15B,C). In summary, these data strongly suggest that lateral crus 1 PCs learn to predict the timing of upcoming rewards shaping preparatory activity in the DN to provide a timed amplification signal to the neocortex.

## Discussion

Our results reveal a key contribution of the cerebellum in the generation of preparatory activity in the neocortex during goal-directed behaviour. The DN - one of the output nuclei of the cerebellum - exhibits preparatory signals prior to reward acquisition that closely resemble those in the motor neocortex during motor preparation (Figure 1 & 2; see also (Murakami et al., 2014; Rickert et al., 2009; Svoboda and Li, 2018). Silencing DN activity by photoactivation of PCs in lateral crus 1 caused a very short-latency decrease (<10 ms) in the activity of the majority of ALM neurons exhibiting preparatory activity (Figure 3). This result is consistent with observations that DN provides direct excitatory input to the motor thalamus (Ichinohe et al., 2000; Kelly and Strick, 2003; Middleton and Strick, 1997; Thach and Jones, 1979), which itself is essential for the generation of persistent activity in the neocortex (Guo et al., 2017; Reinhold et al., 2015). Our results also suggest that preparatory activity in the DN emerges via a transient, learnt decrease in the activity of inhibitory PCs in lateral crus 1 during the period prior to reward acquisition (Figures 4 and 5). Thus, the cerebellum has a specific and fast driving influence on motor cortex activity in anticipation of actions required to acquire rewards. Our results agree with recent work on the role of deep cerebellar nuclei in shaping choice-related activity in ALM as mice perform a tactile discrimination task (Gao et al., 2018), and with human case studies that propose cerebellar contribution to a circuit involving motor thalamus and neocortex in the preparation of self-timed movements (Diener et al., 1989; Purzner et al., 2007).

The activity preceding goal-directed actions has been observed in many brain regions, but its significance is not well understood (Svoboda and Li, 2018). Our study suggests that the preparatory activity we observed in ALM and DN is not directly related to the execution of motor actions. First, although preparatory activity emerged before rewards along with mouse deceleration and anticipatory licks, we found no sign of such activity during deceleration or licking events outside of the reward zone (Figures 1O-R and 2J-M). In fact, preparatory activity only emerged in the rewarded section of the virtual corridor (Figure S3D,G). Those results are corroborated by the GLM classification of neurons exhibiting preparatory activity which found no significant relation between their spike times and lick times or running speed, even though other neurons modulated by licking and running speed were observed in ALM and DN populations (Figures 1F, 2B and S2). Instead, our data argue that preparatory activity reflects a timing signal (Mauritz and Wise, 1986; Roux et al., 2003; Tsujimoto and Sawaguchi, 2005) that predicts the occurrence of the upcoming reward based on elapsed time from learned external cues (Figures S4 and S15), perhaps providing an ‘urgency’ signal that amplifies or primes the emerging motor plan.

The cerebellum is known for its remarkable ability to learn the fine-scale temporal associations between internal and external context and specific actions (Kotani et al., 2003; Mauk and Buonomano, 2004; Medina, 2011; Perrett et al., 1993). We suggest that activity originating within motor-related and sensory areas of the neocortex is conveyed to the cerebellum via the cortico-pontine-mossy fiber pathway where it may be combined with reward prediction signals (Wagner et al., 2017) to adjust the timing of activity in preparation for goal-directed movements (Kunimatsu et al., 2018). The activity of PCs in lateral crus 1 was modulated by running and/or visual flow speed (Figure 5B-G), consistent with this region receiving projections from the visual and hindlimb cortices (Figure S6). However, PC activity decreased significantly more prior to reward than predicted from mouse deceleration profiles during the rest of the trial. This additional activity decrease may result from long-term depression of parallel fibre inputs to PCs (Lisberger et al., 1996) supervised by putative climbing fibre events that occur at reward delivery (Figure 5H-M). Indeed, decreased activity in a large fraction of PCs in advance of reward acquisition is reminiscent of activity profiles resulting from associative learning in cerebellum-dependent tasks such as eyelid conditioning (Freeman et al., 2005; Jirenhed et al., 2007), smooth pursuit (Medina and Lisberger, 2008) and saccadic eye movement tasks (Herzfeld et al., 2018). Additionally, reward time signalling from climbing fibres in lateral crus 1 is consistent with the view that the repertoire of climbing fibre activity extends beyond reporting motor errors and comprises signalling of salient events not directly related to motor performance (Heffley et al., 2018). We hypothesise that the cerebellum learns to predict upcoming rewards to sculpt the timing of preparatory signals generated in the neocortex and maintained in the thalamocortical loop.

The suppressive effects of cerebellar PC photoactivation on ALM activity were predominantly observed in neurons exhibiting preparatory activity prior to reward acquisition, and much less in neurons that responded after reward delivery (Figures 3 and S8). The very short latency suppression suggests the involvement of the tri-synaptic pathway from cerebellar cortex to ALM neocortex (PC-DN-ventral thalamus-ALM) is preferentially influencing a subset of ALM neurons that may be involved in motor planning (Svoboda and Li, 2018). Moreover, the short-latency nature of this effect discards the possibility that the disruption of preparatory activity results from the transient change in mouse motor behaviour which followed PC photoactivation at longer delays.

Following cessation of PC photoactivation the activity in ALM was rapidly reinstated. Because the contralateral ALM has been shown to reinstate preparatory activity at the end of the photoinhibition in the other hemisphere (Li et al., 2016), we examined whether bilateral PC photoactivation would prevent the recovery of ALM preparatory activity (Figure 3) and found that it did not (Figure S10). While the cerebellar output contributes robustly to ALM activity (Fig. 3), it is possible that other cortical regions involved in sensory processing or navigation keep track of the reward context and that this information re-establishes cerebello-cortical activity once this circuit recovers. The basal ganglia have also been shown to process preparatory activity (Kunimatsu et al., 2018; Ohmae et al., 2017) and might therefore contribute a separate subcortical signal for motor preparation.

Given that multiple closed-loop circuits have been identified between the subdivisions of cerebellum and the neocortex (Habas et al., 2009; Kelly and Strick, 2003; Middleton and Strick, 1997; Proville et al., 2014; Ramnani, 2006) we suggest that ALM and the lateral crus 1-DN cerebellar pathway constitutes one such circuit dedicated to the generation of precisely-timed preparatory activity. A recent study has confirmed the existence of a full functional loop between ALM and the cerebellum, involving the DN and FN in maintaining choice-related signals (Gao et al., 2018). This study found that the disruption of DN activity impairs motor preparatory activity in ALM, in keeping with our results, but differences in the effect of manipulating DN and FN activity on ALM choice signals (Gao et al., 2018). Further work will be required to dissect the cerebellar computation giving rise to the FN output involved in motor preparation and how it may complement the role of the lateral crus 1-DN circuit.

More generally, our data add to the growing body of evidence that persistent activity in the neocortex is not a result of recurrent neural interactions within local circuits, but instead requires the coordination of activity across distal brain regions (Gao et al., 2018; Guo et al., 2017; Reinhold et al., 2015; Schmitt et al., 2017; Siegel and Mauk, 2013). The fact that neurons in the deep cerebellar nuclei send excitatory projections to other thalamic regions sub-serving non-motor cortical areas (Kelly and Strick, 2003; Middleton and Strick, 1997; Proville et al., 2014) suggests that they may contribute to the maintenance of persistent neocortical activity during cognitive tasks requiring attention and working memory (Baumann et al., 2014; Gao et al., 2018; Siegel and Mauk, 2013; Sokolov et al., 2017; Strick et al., 2009).

## Acknowledgements

We thank Mateo Velez-Fort and Francesca Greenstreet for help with experiments, Andrei Khilkevich for help with complex spike detection, and David Digregorio, Tom Otis, Marcus Stephenson-Jones and Petr Znamenskiy for constructive ideas and discussions about this work. The research was funded by the Biozentrum core funds, Sainsbury Wellcome Centre core grant (the Gatsby Foundation and Wellcome), ERC Consolidator Grant and SNSF Project grant.

## Author contributions

F.C. performed experiments. F.C. and A.B. analysed the data. All authors wrote the manuscript.

## Methods

### Animal care and housing

All experimental procedures were carried out in accordance with institutional animal welfare guidelines and licensed by the Veterinary Office of the Canton of Basel, Switzerland or under the UK Animals (Scientific Procedures) Act of 1986 (PPL 70/8116) following local ethical approval. For this study we used 35 male C57BL6 mice (supplied by Janvier labs) and 10 mice (7 males, 3 females) from a transgenic cross between Ai32(RCL-ChR2(H134R)/EYFP) and STOCK Tg(Pcp2-cre)1Amc/J lines aged > 60 days postnatal. Animals were housed in a reverse 12:12 hour light/dark cycle and were food-restricted starting a week after surgery with maximum 20% weight loss. Surgical procedures were carried out aseptically on animals subcutaneously injected with atropine (0.1 mg kg^−1^), dexamethasone (2mg kg^−1^), and a general anaesthetic mixed comprising fentanyl (0.05 mg kg^−1^), midazolam (5mg kg^−1^), and medetomidine (0.5mg kg^−1^). Animals were injected an analgesic (buprenorphine, 0.1 mg kg^−1^), and antibiotics (enrofloxacin, 5 mg kg^−1^) at least 15 minutes prior to the end of the surgery and once every day for two days post-surgery. For intrinsic imaging mice were under 1-2% isoflurane anaesthesia. For acute electrophysiological recordings mice were put under 1-2% isoflurane anaesthesia during the craniotomy procedure and allowed to recover for 1-2 hours before recording.

### Behaviour

Mice were trained for 1-2 weeks to run head-fixed on a Styrofoam cylinder in front of two computer monitors placed 22 cm away from the mouse eyes. Mice were trained only once per day with training duration being 15 minutes on the first day, 30 minutes on the second and third days, and 1 hour per day from then-on regardless of the number of trials performed. Running speed was calculated from the tick count of an optical rotary encoder placed on the axis of the wheel with a Teensy USB development board and was fed back as position to a Unity software to display visual flow of a virtual corridor using a MATLAB-based script. A reward delivery spout was positioned under the snout of the mouse from which a drop of soy milk was delivered at a defined position inside the corridor (at 360 cm from start). Licks were detected with a piezo disc sensor placed under the spout and signals were sent to the Teensy USB development board and extracted as digital signals. The virtual corridor was composed of a black and white random dot pattern on a grey background (80 cm long) followed by black and white checkerboard (40 cm long), black and white random triangle pattern on a grey background (80 cm long), vertical black and white grating (40 cm long), black and white random square pattern on a grey background (80 cm long), and a final black and white checkerboard inside which reward was delivered 40 cm from its beginning. The checkerboard pattern was maintained for 2.5 seconds following reward delivery, after which the corridor was reset to the starting position. Mice were initially trained on a short version of the corridor (20, 10, 20, 10, 20 cm length for each visual pattern respectively, and reward position at 90 cm), before extending the corridor to full length. Appearance of the visual patterns inside the virtual corridor was signalled by TCP when the mouse reached the corresponding position in the virtual corridor. In 4/5 mice shown in Figure 3, 3/3 mice from Figure S10 and 3/3 mice from Figure S11, the corridor started at 120 cm distance from start to increase the number of trials.

### Virus and tracer injection

AAV2/1-Ef1a-eGFP-WPRE (30nl, 1.5e^11^ titre) was injected over 15-30 minutes with a *Toohey* Spritzer *Pressure System* (*Toohey Company*) with pulse duration from 5 to 20 milliseconds delivered at 1Hz with pressure between 5 and 15 psi into the left cerebellar crus 1 at the following coordinates: 6 millimetres posterior to Bregma, 3.3 mm Medio lateral, and at a depth of 200 μm. Two weeks after injection mice were euthanized with a dose of pentobarbital (80 mg kg^−1^) and transcardially perfused with 4% paraformaldehyde. Perfused brains were put inside a block of agarose and sliced at 100 μm with a microtome. Slices were then mounted with a mixture of mounting medium and DAPI staining and imaged on a Zeiss LSM700 confocal microscope with a 40X oil objective.

### Intrinsic signal imaging

Mice were anesthetized under 1-2% isoflurane and placed in a stereotaxic frame. A scalp incision was made along the midline of the head and the skull was cleaned and scraped. Two 80 μm tungsten wires (GoodFellow) were inserted inside polyimide tubes (230 μm O.D., 125 μm I.D.) and implanted 300 μm apart into the right primary visual (VisP) and limb motor cortex (lM1) following stereotaxic coordinates (2.7 posterior and 2.5 mm lateral to bregma, 0.25 anterior and 1.5 mm lateral to bregma, respectively) at 800 μm depth from the surface of the brain. Dental cement was added to join the wires to the skull. Neck muscles covering the bone over the cerebellum on the left side were gently detached and cut with fine scissors. The surface of the cerebellum was then carefully cleaned.

Animals were then placed inside a custom-built frame to incline the head and expose the surface of the bone above the cerebellum for imaging with a tandem lens macroscope. Mineral oil was applied to allow imaging through the bone. The mouse was lightly anaesthetized with 0.5-1% isoflurane and the body temperature monitored with a rectal probe and maintained at 37°C. The preparation was illuminated with 700 nm light from an LED source and the imaging plane was focused 300 μm below the skull surface. Images were acquired through a bandpass filter centered at 700 nm with 10 nm bandwidth (Edmund Optics) at 6.25 Hz with a 12-bit CCD camera (1300QF; VDS Vossküller) connected to a frame grabber (PCI-1422; National Instruments).

Tungsten wires were clamped with micro alligator clips and connected to a stimulus isolator (A395; World Precision Instruments). After a 10 s long baseline, trains of 700 μA stimuli were delivered at 6 Hz with pulse duration of 200 μs for 3 s to each cortical area alternatively, followed by a 10 s long recovery phase. Averages of 20 trials were calculated and hemodynamic signals were measured relative to the last 3 s before stimulation (∆F/F_0_). Location of tungsten electrodes inside the neocortex were confirmed post-hoc with DiI labelling of the tracts.

### In vivo extracellular electrophysiology

Mice were anaesthetized according to the surgical procedure described in the animal care and housing section and placed into a stereotaxic frame. The skin over the skull was incised along the midline and the skull was cleaned and scrapped. A headplate was then attached to the skull in front of the cerebellum using Super Bond dental cement (Super-Bond C&B). For cerebellar recordings the neck muscles covering the bone were gently detached and cut with fine scissors on the left side. The surface of the skull over the cerebellum was then cleaned, a small piece of curved plastic was glued to the base of the exposed skull to support a well attached to the headplate and built up with dental cement and Tetric EvoFlow (Ivoclar Vivadent). The well was then filled with Kwik-Cast sealant (World Precision Instruments). For the simultaneous recordings in cerebellum and ALM, a small additional well was built around stereotaxically-defined coordinates for the right ALM (2.5 mm anterior and 1.5 mm lateral to bregma).

On the day of the recording mice were anaesthetized under 1-2% isoflurane and small craniotomies (1mm diameter) were made above left lateral crus 1 (6 mm posterior and 3.3 mm lateral to bregma), left dentate nucleus (6 mm posterior, and 2.25 mm lateral to bregma), and/or right ALM (2.5 mm anterior and 1.5 mm lateral to bregma). Mice recovered from surgery for 1-2 hours before recording. Mice were then head-fixed over a Styrofoam cylinder. The well(s) around the craniotomy(ies) were filled with cortex buffer containing (in mM) 125 NaCl, 5 KCl, 10 Glucose monohydrate, 10 Hepes, 2 MgSO_4_ heptahydrate, 2 CaCl_2_ adjusted to pH 7.4 with NaOH. A silver wire was placed in the bath for referencing. Extracellular spikes were recorded using NeuroNexus silicon probes (A2×32-5mm-25-200-177-A64). The 64- or 128-channel voltages were acquired through amplifier boards (RHD2132, Intant Technologies) at 30 kHz per channel, serially digitized and send to an Open Ephys acquisition board via a SPI interface cable. Mice were recorded up to 90 minutes only once regardless of the number of trials performed (ranges from 36 to 333, 137 +/− 86 mean and SD) except for 1/5 mouse from Figure 3 and 2/6 mice from Figures S10 and 11 that were recorded over 2 consecutive days, in which case the cement wells around the craniotomies were filled with Kwik-Sil (World Precision Instruments) and sealed with Tetric EvoFlow after the first recording session.

### Photoactivation

A 200 μm diameter optical fibre was placed on top of the surface of left lateral crus 1 using a manual micromanipulator. Light was delivered by a 100 mW 473 nm laser (CNI, MBL-III-473) triggered by a Pulse Pal pulse train generator (Open Ephys) using 1-second long square pulses (*n* = 5 recordings, 5 mice). In one additional recording (data included in Figure 3 L-Q) the photoactivation period lasted 2 seconds. To prevent mice from seeing the laser light, a masking 470 nm light from a fibre-coupled LED (Thorlabs) was placed in front of the connector between the patch cable and the optical fibre and turned on during the whole recording session. Mice were also trained in the presence of LED light. Black varnish was painted over the cement well surrounding the craniotomy and black tape was wrapped around the connection between the patch cable and the optical fibre. One-second square light pulses (5 to 10 mW) were randomly delivered in 40 % of trials (at least 10 trials, *n* = 38 +/−28, mean and SD from 5 mice). Control trials from mice that experienced photoactivation were not included in Figures 1 and 2 to avoid confounding effects such as plasticity-induced change in neuronal activity. For bilateral crus 1 photoactivation, a second 200 μm diameter optical fibre was placed over the right crus 1 and was coupled to a second 80 mW Stradus 473-80 nm laser (Vortran Laser Technology, Inc). The light pulses were set as square onsets, continuous voltage for 700 ms (1 to 4.5 mW), and ramp offsets in the last 300 ms to limit the rebound of activity in DCN neurons. Unilateral (left lateral crus 1 only) and bilateral stimulations occurred randomly in 40% of trials (at least 10 trials each, *n* = 24 +/−17 and 24 +/− 16, mean and SD from 3 mice). The same protocol was used for lateral crus 1 vs lobule IV/V photoactivation (*n* = 26 +/−10 and 33 +/−32 trials, mean and SD from 3 mice).

### Electrophysiology data analysis

Spikes were sorted with Kilosort (https://github.com/cortex-lab/Kilosort) using procedures previously described (Pachitariu et al., 2016). Briefly, the extracellular voltage recordings were high-pass filtered at 300 Hz, the effect of recording artifacts and correlated noise across channels were reduced using common average referencing and data whitening. Putative spikes were detected using an amplitude threshold (4 s.d. of baseline) over the filtered voltage trace and matched to template waveforms. The firing rate for each unit was estimated by convolving a Gaussian kernel with spike times, σ was chosen according to the median inter-spike interval of each individual unit.

For population scatter plots (Figures 1, 2, 3 and 4) and averaging across neuronal activities grouped by type we used the z-score of firing rates.

The cross-correlogram between each PC and DN neuron simultaneously recorded (n = 1855 pairs, 3 mice) was computed with a bin of 1 ms (Figure 4M). A correlogram was considered as modulated if at least two consecutive bins in the 10 ms following the Purkinje cell spike were above 3 SD of the baseline computed in the [-50, −10] window. For all these pairs (46/1855) the cross-correlogram was z-scored by the mean/SD of this baseline and all z-scored correlograms were averaged. The strength of connection was measured as the average of the z-scored correlograms between 1 and 7 ms (Figure S14) and pairs were split between excited (n = 6) and inhibited (n = 40) based on the sign of this average. For Figure S14D, the response profiles of PCs and DN neurons around reward time were computed and z-scored using a baseline taken between −10 and −5 seconds. For inhibited pairs, the spearman correlation coefficient of these response profiles in the 5 seconds before reward was correlated to the strength of connection using a linear regression model (matlab fitlm, blue line Figure S14E). Shaded area indicates the 95% confidence intervals.

On longer time scale, task modulation of the neurons entrains instabilities of the firing rate that might produce spurious covariance between comodulated pairs. To assess the relation between PC activity and DN neuron activity on these time scales we used two equivalent methods. In Figure 4I, the cross-covariance between firing rates of PC and DN pairs was corrected for correlated firing resulting from stimulus effects by subtracting the cross-covariance between shuffled trials and was then normalized by the variance of the firing rates. In Figure 4N, the cross-correlogram between each pair was first calculated on each trial in the last 10 second before the reward (CC_raw_). We then computed the cross-correlogram for the same pair but using trial n and n+1 (CC_shuffled_). The shuffled corrected correlogram was then defined as (CC_raw_ – CC_shuffled_) / sqrt(CC_shuffled_) and averaged across pairs.

ALM neurons were considered modulated by cerebellar photoactivation if the average firing rate in the second following the onset of photostimulation was significantly (rank-sum, alpha of 0.05) different from the average firing rate during the same window in control trials. We classified them as excited/inhibited if the control response was lower/higher than that during photoactivation trials. Average firing rate of the population in the same 1 s window were compared between control and photoactivation condition using signed-rank test (alpha 0.05). Z-scored activity profiles were obtained for each neuron by subtracting the average firing rate of the neuron across the whole recording from the neuron average activity profile in Hz and dividing it by the SD of the firing rate. The z-scored activity profiles were then averaged together to generate the population activity profile (Figure 3 H-K). The onset of inhibition (Figure 3Q, Figure S7D,E, Figure S10C and Figure S11D) was measured as the first 2 ms bin after 0 where the cross-correlogram was below 2 SD of a baseline measured in the preceding 50 ms. For figure S8, type by type comparisons (Figure S8A,B and G) were done with Wilcoxon ranksum test applying Bonferonni correction, leading to an alpha of 0.0083 for Figure S8A,B and 0.0125 for Figure S8G. To assess the link between control firing rate, ramp size and photoactivation effect (as defined in Figure S8H), we did a simple linear regression model, photoactivation effect = \alpha ramp size + \beta control firing rate \gamma. In this regression, only \alpha was significantly different from 0 (\alpha = −0.79, p = 1.26e-6, \beta = 0.03, p = 0.76 and \gamma = 2.04, p = 0.08).

### Classification of cerebellar cortex and ALM units

Cerebellar cortex units (from crus 1 and crus 2 recordings) were classified as putative Purkinje cells (PCs) by fitting their interspike intervals distributions (Figure S12B) with a lognormal function. The distribution of means and standard deviations from the log normal fits were then clustered in 2 groups using k-means clustering (Figure S12C). Units belonging to the group with higher ISIs mean and SD were considered as non-PCs. This group likely comprises inhibitory interneurons found in the molecular layer (Blot et al., 2016; Gao et al., 2012) or in the granule cell layer (Golgi cells) (Gao et al., 2012), but might also contain PCs with low firing rates or PCs whom recording quality deteriorated during the recording session. We nonetheless adopted this conservative measure to avoid contaminating of PCs sample with other inhibitory interneurons.

Accurate classification of units as PCs was confirmed by detecting climbing fibre-generated complex spikes (CSs, Figure S12E). For every single unit, we first identified putative PCs by applying a 15 Hz threshold to the baseline firing rate. Next, like previous studies (Blot et al., 2016; Khilkevich et al., 2018; de Solages et al., 2008), we applied analyses that aimed to separate simple and complex spikes based on consistent differences in their typical waveforms. While the exact waveform of complex spikes can vary substantially, a common feature is a presence of positive deflection following the spike peak, to which we will refer as “late peak”. For all waveforms belonging to a single unit, we calculated the distribution of late peak values within 3 ms following the spike peak time. Identified waveforms with late peak values larger than three median absolute deviations (MAD) of the median late peak value were selected. The mean profile of these waveforms was used as a proxy for putative complex spike waveform. The mean profile of the rest of waveforms resulted in a simple spike waveform. For every spike, we calculated Spearman correlation coefficients between corresponding waveform and the mean waveforms of complex and simple spikes. The combination of: positive correlation with mean complex spike waveform and; a positive difference between correlations with mean complex and simple spike waveforms was used to separate putative complex spikes from simple spikes. Complex spikes were confirmed if their cross-correlogram with simple spikes from the same unit (Figure S12D) exhibited tens of ms-long pauses in SS firing (Figure S11F). In crus 1 (19/72) and in crus 2 (13/56) units classified as PCs exhibited CSs but none classified as non-PCs.

In crus 1 we also confirmed correct unit classifications as PCs if those units had significantly modulated cross-correlograms with ms-latency through with DN units (Figure 4L). Ten units classified as PCs exhibited such cross-correlograms. Three units classified as non-PCs had similar cross-correlograms with DN units and were integrated in the PCs group. In Figures 4,5 and S12 only putative PCs (72/89 of all units for crus1 and 56/84 for crus 2) are included.

Units were classified as ‘fat spikes’ (Figure S12G) (Gao et al., 2012) if the full width at half maximum of their spike waveform exceeded 500 ms (Figure S12H). Fat spikes units were considered to exhibit significant increase in spiking probability if the number of spikes in the last 500 ms before reward were significantly lower than in the first 500 ms after reward across trials (p < 0.05, Wilcoxon Rank-sum test) (Figure 5I-M).

ALM units were classified as putative pyramidal neurons or fast-spiking interneurons based on spike width as described in (Guo et al., 2014) and only putative pyramidal neurons were analysed.

### Generalized linear model

We used neuroGLM (https://github.com/pillowlab/neuroGLM.git) to classify neuronal responses with models obtained from linear regression between external covariates and spike trains in single trials. Spike trains were discretized into 1 ms bins and each external event was represented as follows: running speed was added as a continuous variable. Reward times, lick times, and visual cue times were represented as boxcar functions convolved with smooth temporal basis functions defined by raised cosine bumps separated by π/2 radians (25 ms). The resulting basis functions covered a −4 to 2 s window centred on reward time, and −2 to 2 s windows for lick and visual cue times. We then computed Poisson regression between spike trains and the basis functions and running speed. The resulting weight vectors were then convolved with event times and linearly fitted with the spike times peri-stimulus time histograms smoothed with a 25 (for lick times) or 50 ms Gaussian (for reward times and running speed) to compute the coefficient of determination for each trial (Figure S2). We divided the fit between reward times model and firing rates in two time windows: −4 to 0 s and 0 to 2s relative to reward time to differentiate between pre- and post-reward neuronal activity (Figure S2A-F). Fits with mean coefficient of determination across trials exceeding 0.17 were selected to classify units (Figure S2K).

### Linear modelling of Purkinje cells firing rate from running speed

For each mouse, the running speed was low-pass filtered at 100 Hz and discretized in bins of 5 cm/s from the minimum to the maximum values rounded to the nearest integers. The times of mouse running speed were then sorted according to which bin the running speed fell into and the firing rate of PCs at those times was then extracted and averaged. The resulting PCs firing rates to running speed tuning curves were fit using the MATLAB smoothing spline function. PCs activity models were obtained by converting the running speed from the filtered trace to the corresponding firing rates on the tuning curves fits. For this analysis we included only the PCs that exhibited decreases in firing rates preceding reward times satisfying the following condition: mean(FR[-4,-3]s) > mean(FR[-2,0]s)+2*SD(FR[-4,-3]s), where time in square brackets is relative to reward times.

The averaged modelled firing rates were then subtracted from the averaged PCs firing rates around deceleration events outside of the reward zone ([-2,0]s, mean(DecelModelFR) – mean(DecelPcFR)) and around reward times ([-2,0]s, mean(RewardModelFR) – mean(RewardPcFR)). In most cases the number of reward times exceed that of deceleration events and the former were then grouped in blocks (i) containing the same number of trials than in the latter. Models were considered to overestimate the PCs firing rates around reward times if mean(DecelModelFR)-mean(DecelPcFr) > mean(RewardModelFR)-mean(RewardPcFR(i)) in at least 80% of cases. The plot in Figure 5G shows the mean values across all reward time blocks for the ‘Reward’ condition (right column).

### Data availability

The data that support the findings of this study are available from the corresponding author upon reasonable request.

---

**Figure S1.**
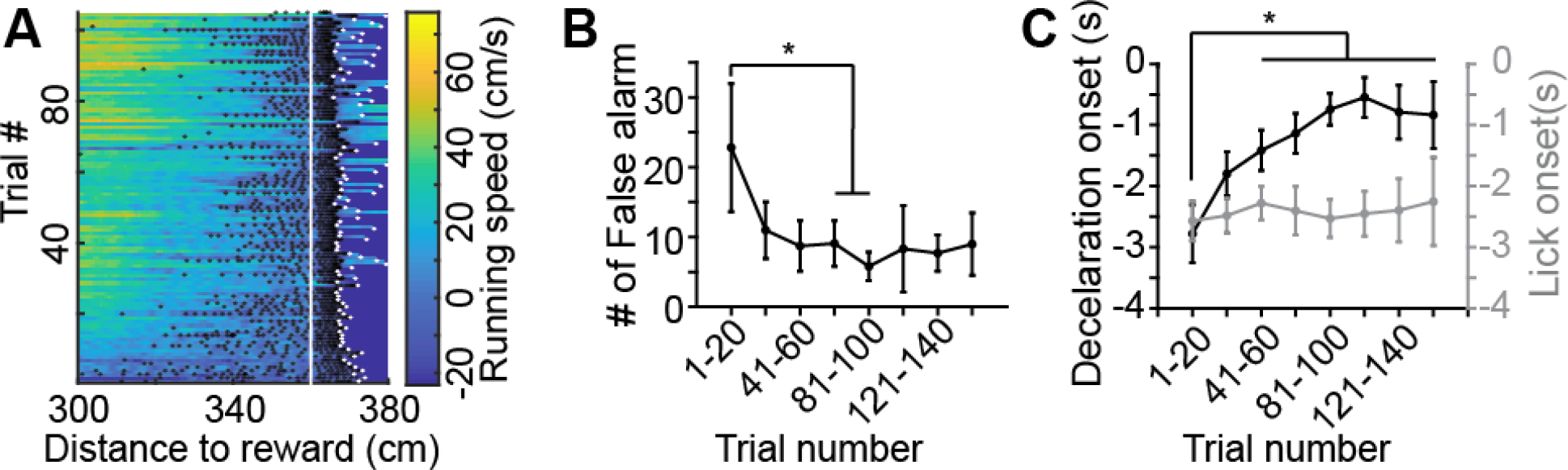
Refinement of motor behaviour in anticipation of reward (related to Figure 1) (A) Summary of behavior inside the rewarded section of the virtual corridor for an example recording session of a trained mouse (including all trials). Running speed over distance is color-coded. Black and white dots represent individual lick times and end of trial, respectively. Reward position is indicated by the vertical white line. (B) Number of false alarm licks averaged in chunks of 20 trials over the course of recording sessions across all mice (*n* = 21). Error bars are standard deviations. Multiple comparison test after a Kruskal-Wallis test shows significant decrease of false alarm licks over 61-80 trials. (C) Summary plots of deceleration and lick rate onsets relative to reward time (respectively, onset of 20% decrease and increase of Z-score values) averaged in chunks of 20 trials across recording sessions. Error bars are standard deviations. Time between deceleration onset and reward delivery significantly decreased over 41-60 trials (multiple comparison test after a Kruskal-Wallis test).

**Figure S2.**
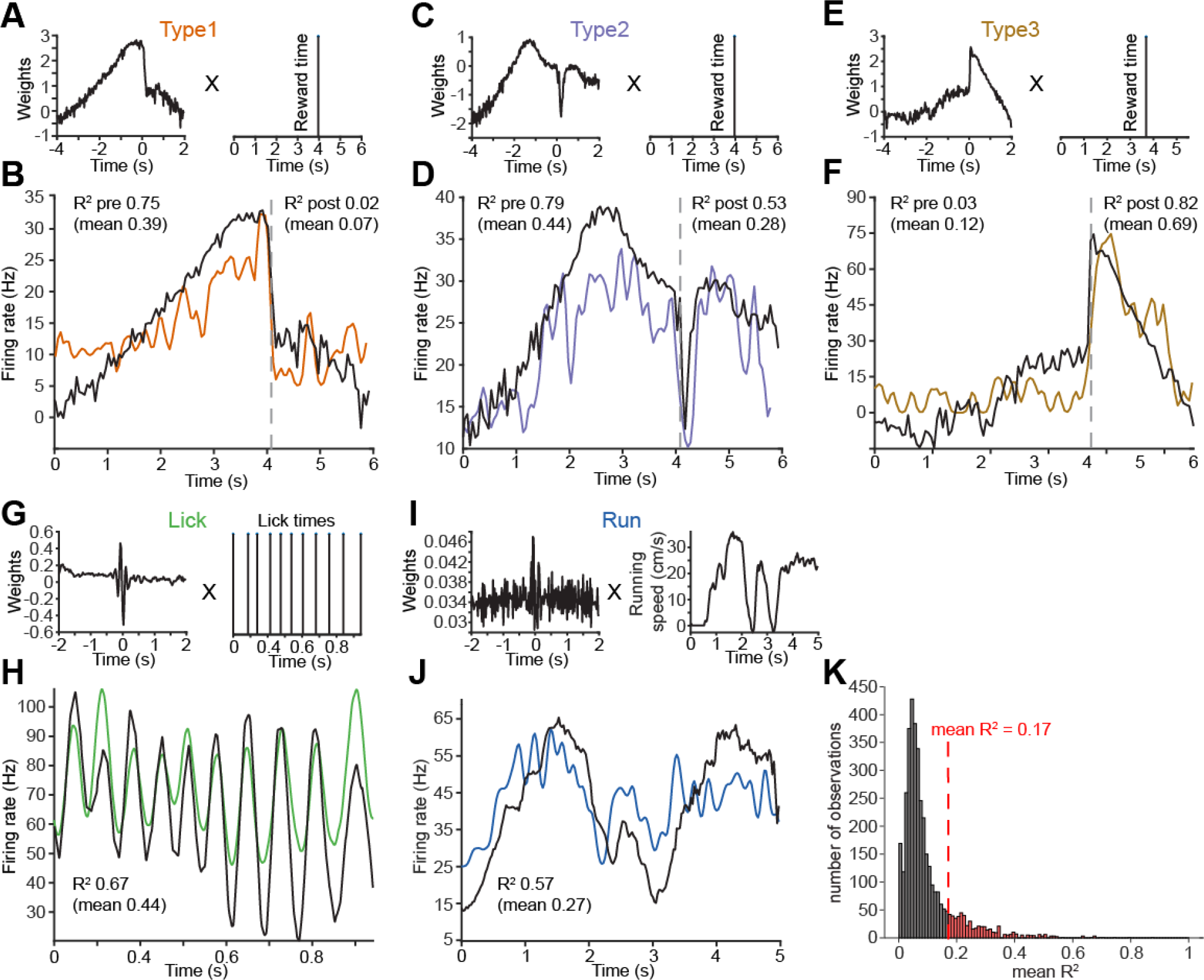
Unit classification using generalized linear model single-trial predictions from external covariates (related to Figures 1,2 and 3) (A) The weight vector (left) resulting from linear regression of spike times and reward times for a unit classified as type 1 is convolved with single trial events (reward time, right) to model neuronal activity. The goodness of fit of GLM model (black trace) to neuronal firing rates (color-coded trace) are estimated over single trials. The coefficient of determination value for the example trial is written in the panel, the mean coefficient of determination across trials is given in parentheses. For reward times-based models the fit was evaluated over 2 time windows: −4 to 0, and 0 to 2 s around reward times, as indicated by the grey dashed line, to distinguish pre-reward only (type 1), pre and post-reward (type 2), and post-reward only (type 3) neuronal activity. Coefficient of determination values for pre- and post-reward fits are given on the left side and right side of the traces respectively (B, D, F). (C, D) Same as (A,B) for a unit classified as type 2. (E, F) Same as (A,B) for a unit classified as type 3. (G, H) Same as (A,B) for a unit with activity best predicted by lick times. (I, J) Same as (A,B) for a unit with activity best predicted by running speed. (K) Distribution of mean coefficient of determination values between all units and GLM models obtained from each external covariate (reward times, lick times, and running speed). A threshold criterion for significant goodness of fit (mean R^2^ > 0.17, red dashed line) was obtained by performing k-means clustering on the R^2^ distribution with 2 clusters.

**Figure S3.**
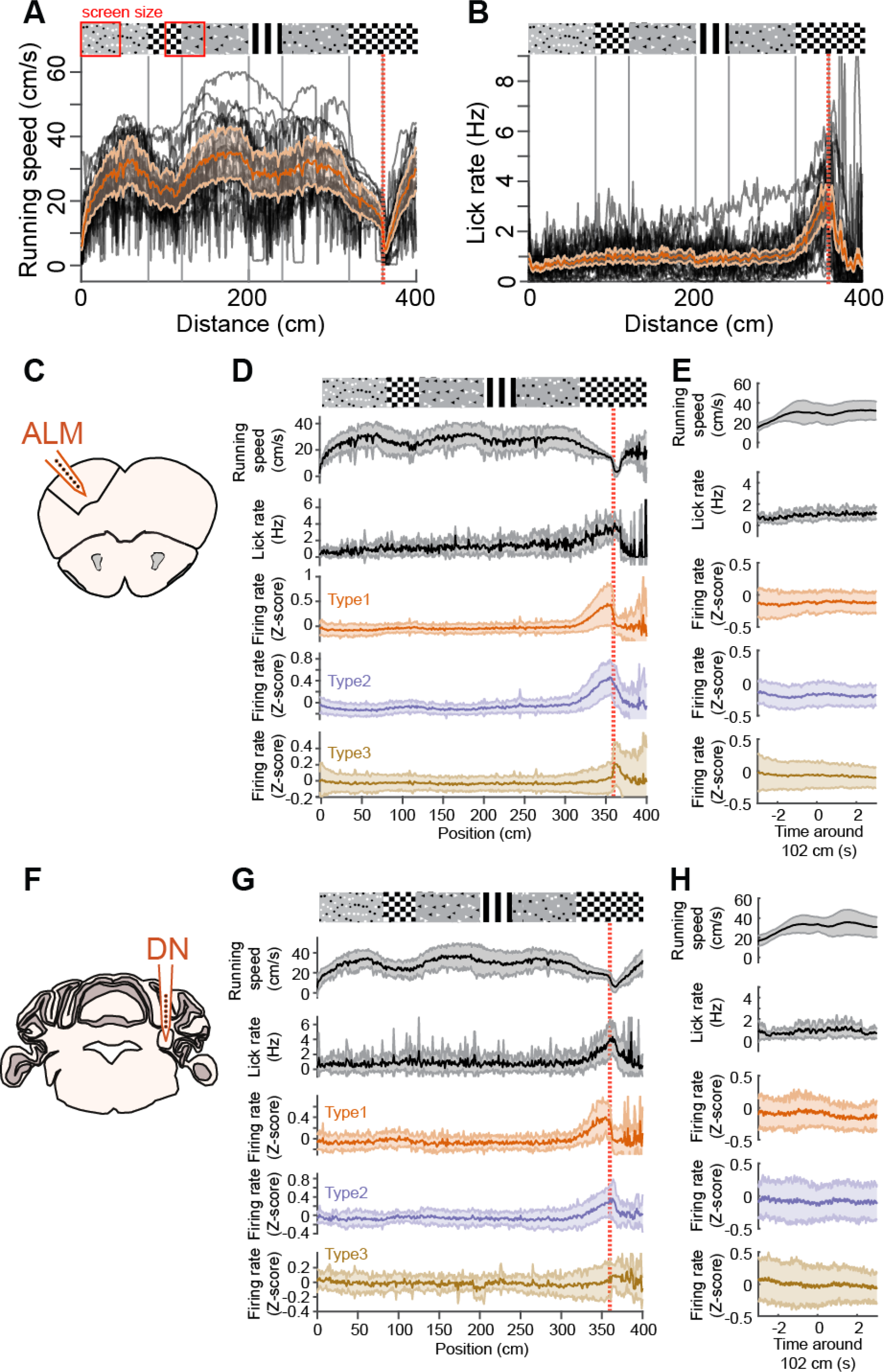
Type 1-3 ALM and DN neurons are selectively active around reward position (related to Figure 1 and Figure 2) (A) Running speed profiles for all mice (black curves, 21 expert mice comprising the 6 ALM and 15 DN recordings shown below) and population average (orange trace, shading is SD) as a function of position inside the virtual corridor. The reward position is indicated by the red vertical dotted line. The visual patterns displayed on the corridor walls shown above the running traces are aligned to the position at which they fully appear in the field of view of the mice (i.e. when they reach the back edge of the monitors). The full field of view of the mouse which corresponds to 47 cm (monitor width) is indicated by the red square box around the visual patterns. Note the consistent deceleration of mice around the 1^st^ non-rewarded checkerboard and to a lesser extent around vertical gratings. The trough of mouse speed around the 1^st^ checkerboard occurs around 102 cm, when the mouse is halfway out of the checkerboard as indicated by the second red box. (B) Same as (A) but for lick rate. (C) Schematic showing recording location in the anterolateral motor cortex (ALM). (D) Type 1-3 ALM neurons (line is mean and shaded area is SD from 37, 29, and 88 neurons respectively, recorded from 6 mice) are mostly active around reward position (vertical red dotted line). Note the slight increase in neuronal activity around the 1^st^ non-rewarded checkboard at which position mice decelerate. (E) Mean (line) and SD (shaded area) of running speed, lick rate, and type 1-3 ALM neurons activity around the time of mice deceleration trough occurring after the appearance of the 1^st^ checkerboard (at 102 centimetres from the start of the corridor). (F) Schematic showing recording location in the Dentate nucleus (DN). (G) same as (D) for type 1-3 DN neurons (20, 32, and 57 neurons respectively, recorded from 15 mice). (H) same as (E) for type 1-3 DN neurons.

**Figure S4.**
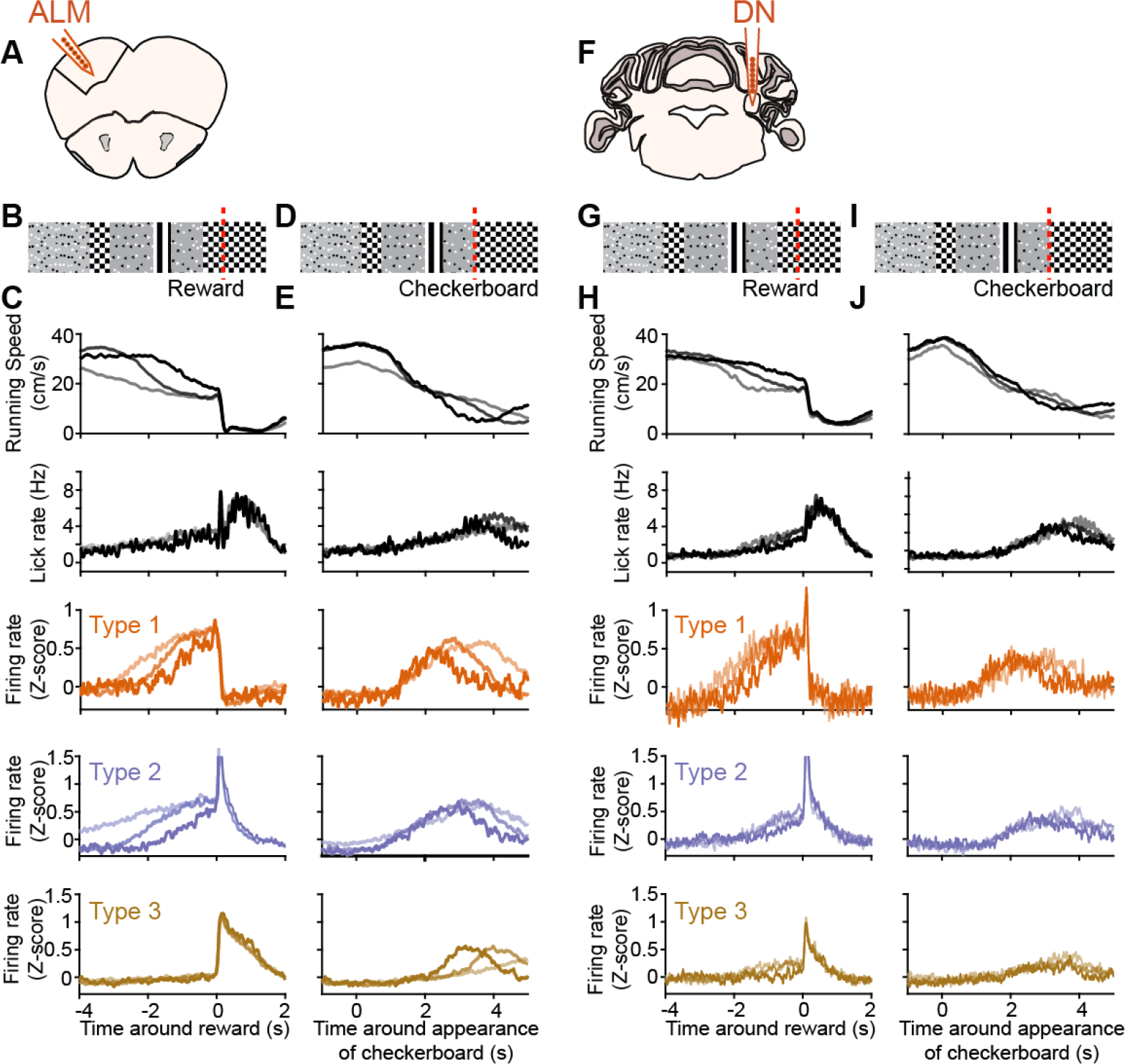
Dynamics of ALM and DN preparatory activity are consistent with encoding time between cue and reward delivery (related to Figure 1 and Figure 2) (A) Schematic showing recording location in the anterolateral motor cortex (ALM). (B) Schematic of virtual corridor showing the location of reward delivery (red dotted line). (C) From top to bottom, mean running speed, lick rate, and type 1-3 ALM neuronal firing rate (37, 29, and 88 neurons respectively) binned according to the time at which mouse speed dipped under 20 cm/s in the 4 s before reward (1^st^ group : times below the 33^rd^ percentile of the distribution; 2^nd^ group: times between the 33^rd^ and 66^th^ percentile; 3^rd^ group: times above the 66^th^; *n* = 6 mice). (D) Same as in (B), with the red line denoting the appearance of the rewarded checkerboard. (E) same as (C) aligned to appearance of the rewarded checkerboard. Note the substantial decrease in onset jitter compared to reward-aligned traces (C) for behavioral variables and type 1-2 neurons. (F) Schematic showing recording location in the Dentate nucleus (DN). (G) Same as (B). (H) Same as (C) for DN recordings (20, 32, and 57 type 1-3 neurons respectively, recorded from 15 mice). Note that only type 1 neurons show substantial jitter in onset of ramping. (I) Same as (D). (J) Same as (E) for DN recordings.

**Figure S5.**
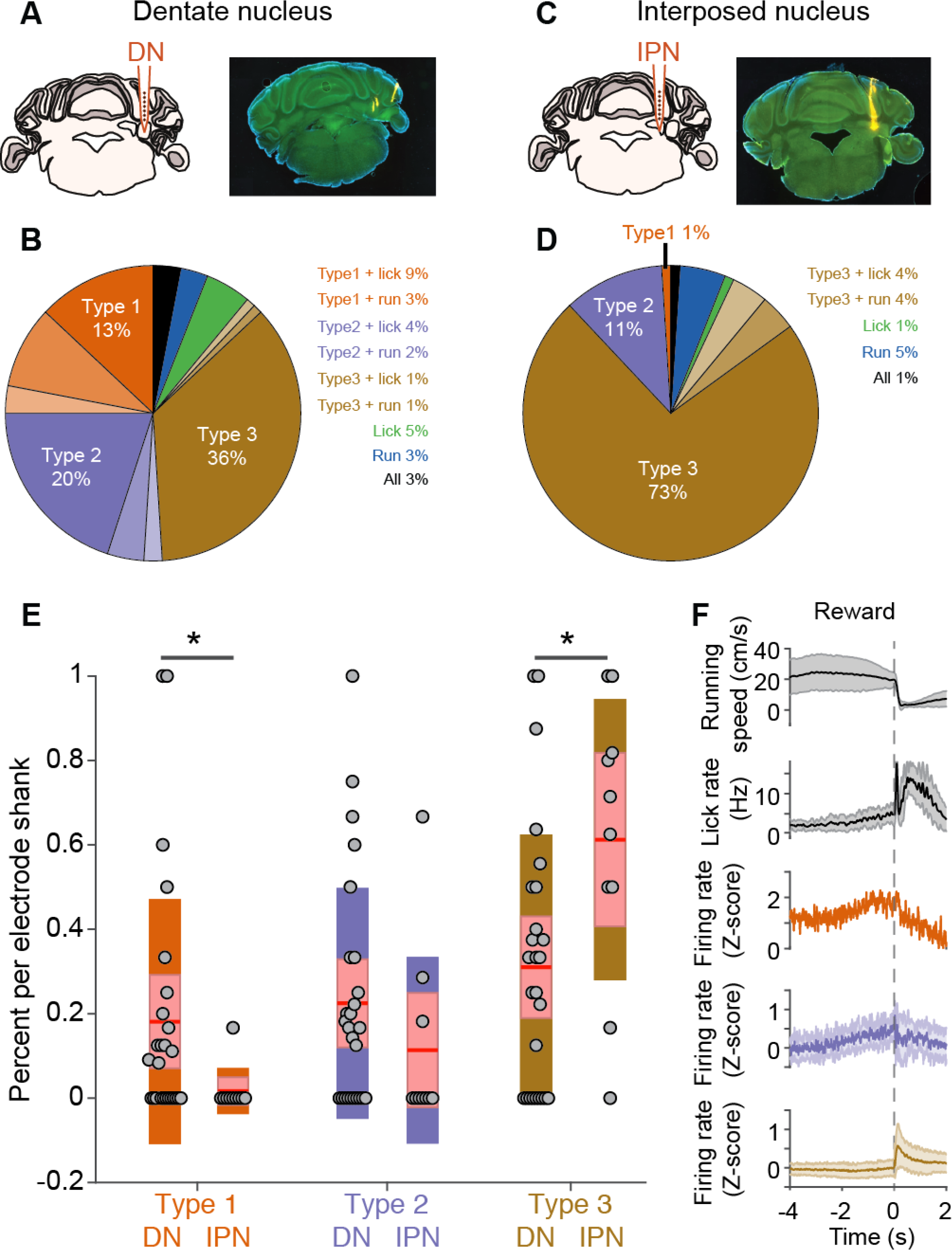
Preparatory activity is significantly more present in the dentate than in the interposed nucleus (related to Figure 2) (A) Left: Schematic of recording location in the Dentate nucleus (DN). Right: Coronal brain slice from a DN recording (yellow traces are from DiI staining on recording electrode shanks, tract in lateral crus 1 is from simultaneous recording in the cerebellar cortex). (B) Summary DN neuron classification. (C) Same as (A) but showing recording location in the more medial interposed nucleus (IPN). (D) Summary IPN neuron classification. (E) Summary plot of percentage of type 1-3 neurons amongst classified units per electrode shank for DN (left columns) and IPN (right columns). Proportion of type 1 neurons is significantly lower in IPN (*n* = 1/74, 5 mice) than in DN (*n* = 20/170, 15 mice; p = 0.024, Wilcoxon Rank-Sum test). There is no significant difference in the proportion of type 2 neurons between IPN (*n* = 8) and DN (*n* = 32; p = 0.16, Wilcoxon Rank-Sum test). Proportion of type 3 neurons is significantly higher in IPN (*n* = 54) than in DN (*n* = 57; p = 0.02, Wilcoxon Rank-Sum test). (F) From top to bottom, average (line) and SD (shaded area) running speed, lick rate, and type 1-3 IPN neuronal activity centered on reward (grey dotted line).

**Figure S6.**
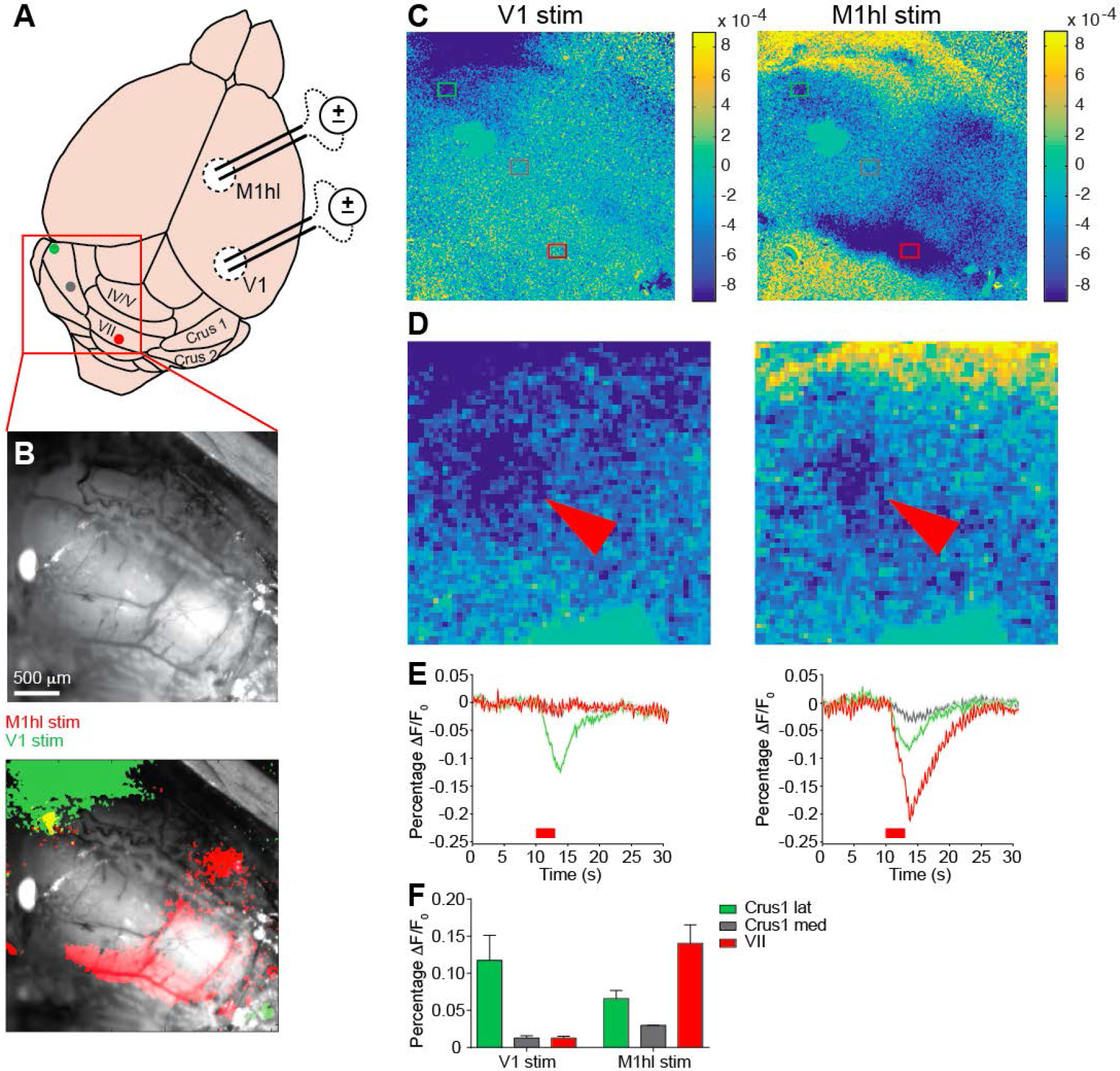
Mapping visuomotor cerebellum (related to Figure 3) (A) Schematic of electrical stimulation for identification of a cerebellar region activated by inputs from primary visual cortex (V1) and hind limb-related motor cortex (M1hl). Coloured dots correspond to regions from where hemodynamic signals were measured in (C-E): lateral crus 1, medial crus 1 and lobule VII. (B) Top: wide-field image of the cerebellar surface. Bottom: same image overlaid with 20-trials of hemodynamic signals averaged across sessions (showing only peak decrease from baseline) for electrical stimulation of V1 (green) and M1hl (red) for an example recording. Note that the large haemodynamic signal induced by V1 stimulation outside of the cerebellum probably comes from the cerebral venous sinus. (C) Wide-field image of 20-trials average hemodynamic signals color-coded according to percentage change of infrared reflectance from baseline for V1 (left) and M1hl stimulation (right). Coloured rectangles indicate the areas from which the signals were measured from. (D) Zoom on the lateral crus 1 zone from the wide-field image in (C). Red arrowheads indicate region of max hemodynamic response. (E) Mean response time courses after V1 (left) and M1hl stimulation (right) color-coded according to the sampled areas in (C). ∆F/F0, normalized change in reflectance. The timing of electrical stimulation is indicated by the red bar below the traces. (F) Summary of peak hemodynamic response values after V1 (left) and M1hl (right) stimulations for each cerebellar area (*n* = 4 mice).

**Figure S7.**
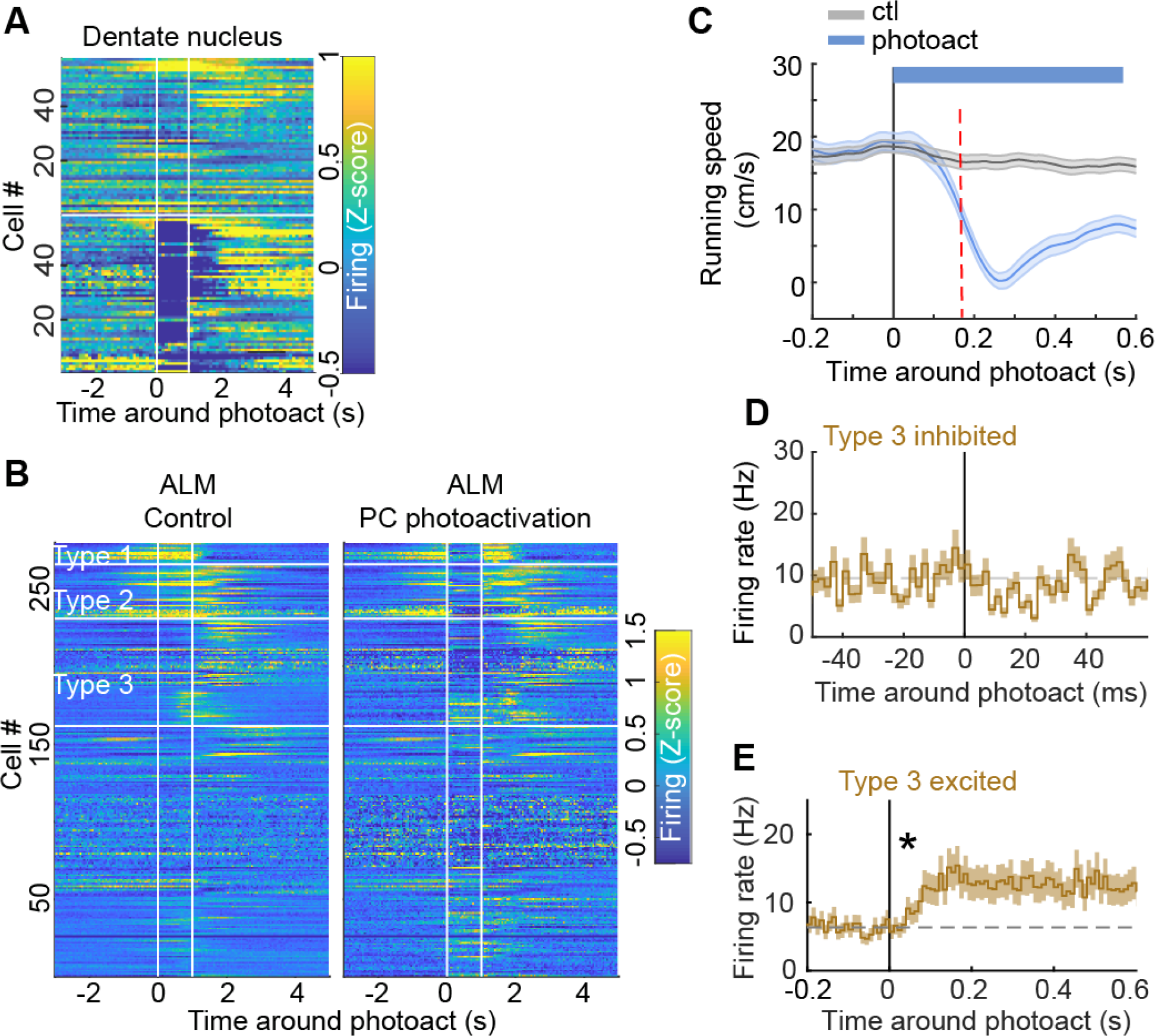
Effects of cerebellar Purkinje cells photoactivation on DN and ALM neuronal populations and on mouse locomotion (related to Figure 3) (A) Purkinje cell (PC) photoactivation efficiently silenced most dentate nucleus (DN) neurons (one line per neuron). Upper half: control trials. Lower half: photoactivation trials. (B) PC photoactivation efficiently suppressed activity in most type 1 and type 2 ALM neurons (one line per neuron). Purkinje cell photoactivation (right) compared to control trials (left). The effects of photoactivation on type 3 and unclassified ALM neurons were more diverse. (C) PC photoactivation (‘photoact’) induced sudden mouse deceleration. The time of significant deviation from running speed in control (‘ctl’) trials is indicated by the red dotted line at 170 milliseconds (running speed significantly lower than control speed minus one standard deviation, calculated from 10 ms bins, p < 0.005, Wilcoxon Rank-Sum test). (D) Average response profiles of firing rate type 3 ALM neurons with significant reduction in activity compared to control trials, during the first 60 milliseconds following PC photostimulation onset (vertical line). Shaded area represents SEM. (E) Same as in (D) with 10 ms binning for type 3 ALM neurons which activity significantly increases after PC photoactivation (star: first significant bin at 40 millisecond). Shaded area represents SEM.

**Figure S8.**
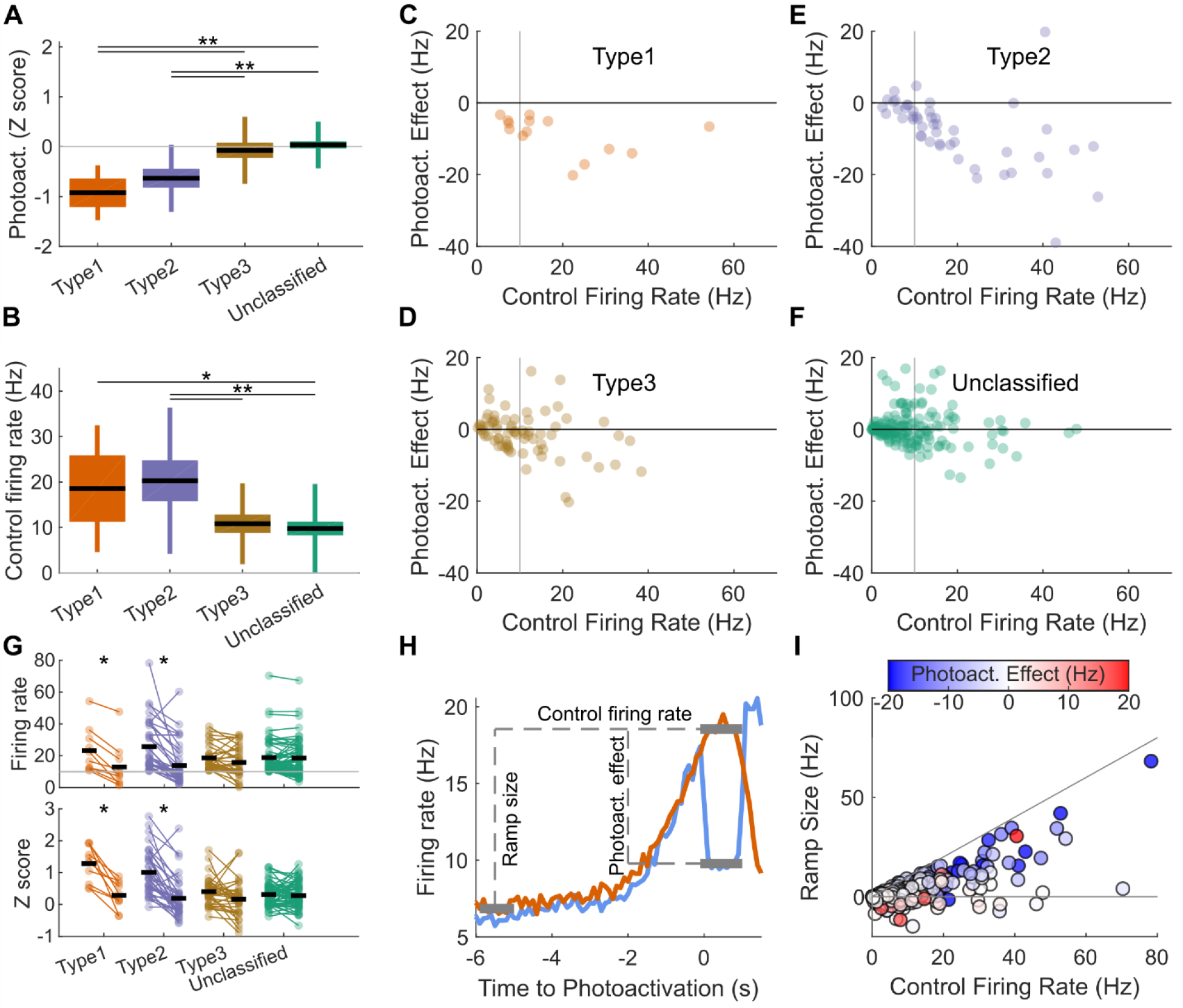
Photoactivation effect is not explained by firing rate (related to Figure 3) (A) Type 1 and type 2 neurons were significantly more inhibited by photoactivation than type 3 and unclassified neurons. (B) Type 1 and type 2 neurons had higher firing rate in control trials. (C-F) The effect of photoactivation did not seem to be explained by this difference in firing rate as type 1 (C) and type 2 neurons (E) were mostly inhibited regardless of firing rate and inhibited type 3 (D) and unclassified neurons (F) had similar firing rates to that of unmodulated neurons. (G) When selecting only neurons firing at more than 10 Hz in control condition (vertical grey line in C-F), type 1 and type 2 neurons were significantly inhibited while the firing rate of type 3 and unclassified neurons was not significantly affected (see Methods). (H-I) The photoactivation effect (blue trace) was correlated with the size of the ramp (orange trace) but not with the control firing rate (see Methods). * indicates p < 0.005, ** p < 0.0001. In (A) and (B), box plots represent the 95 % confidence intervals and SD, black line is the median. In (A-G), *n* = 14, 49, 74, 165 respectively for Type 1, 2, 3 and unclassified neurons. In (I), *n* = 302 neurons, including all types.

**Figure S9.**
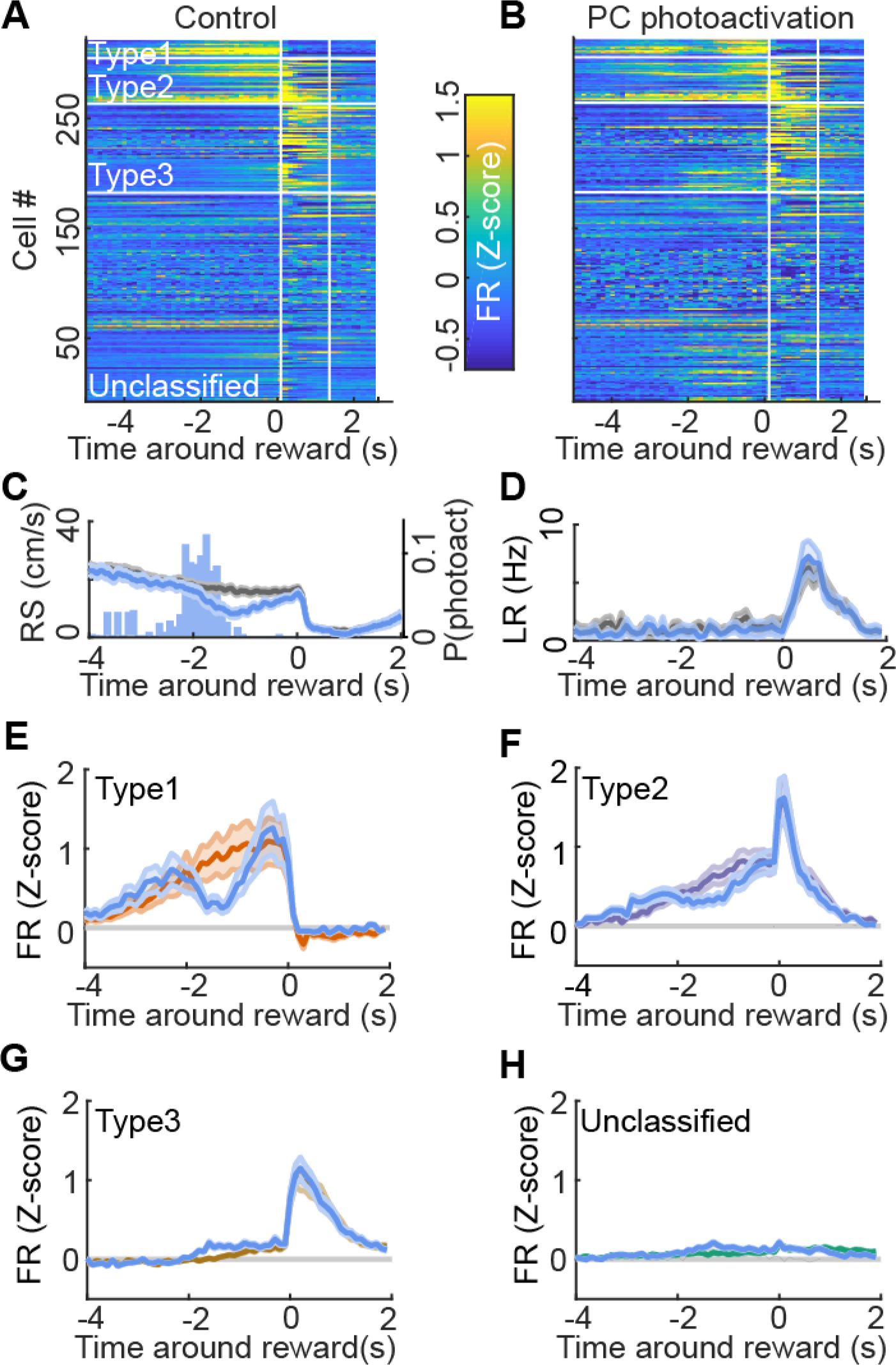
Effects of cerebellar photoactivation on ALM activity aligned to reward time (related to Figure 3) (A,B) Response profiles colour-coded by firing rate (z-scored) for all ALM neurons sorted by cell type in control condition (A) and with photoactivation of Purkinje cells (PC, B) aligned on time of reward delivery. The photoactivation started 20 cm in advance of the reward position, corresponding to about 2 s before reward time (see photoactivation probability distribution in panel (C)), and lasted 1 s. (C,D) Average profiles around reward time of stimulation in photoactivation trials (blue lines) and control trials (grey lines) for running speed (RS, C) and lick rate (LR, D) and distribution of photoactivation probability (P(photoact)) in reward trials (C, blue histogram). (E-H) Firing rate profiles aligned to reward time for type 1 (E), type 2 (F), type 3 (G) and unclassified ALM neurons (H) during photoactivation (blue traces) and control trials (coloured traces). Shaded areas in (C-H) represent SD.

**Figure S10.**
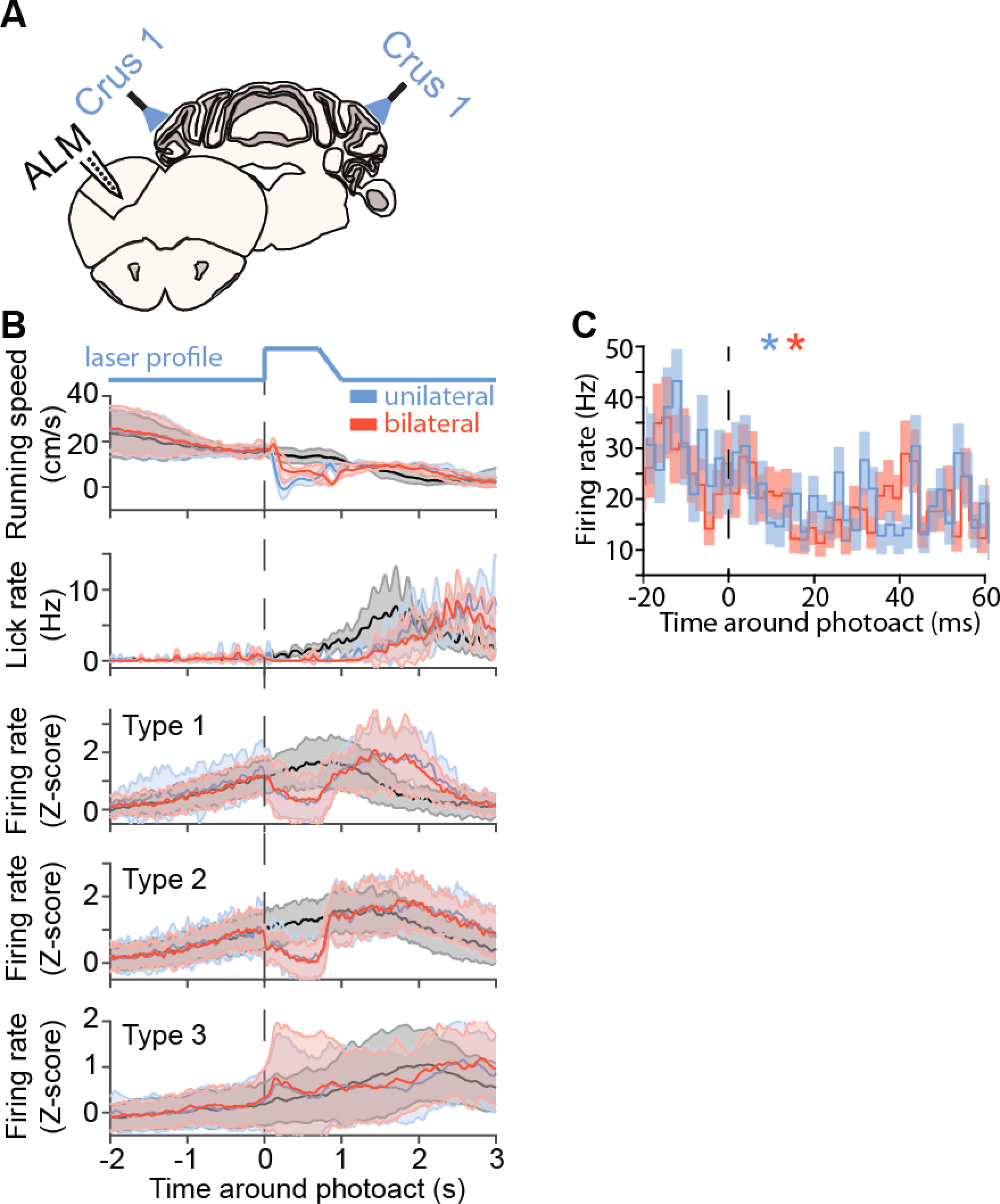
Comparison of the effect of unilateral and bilateral crus 1 photostimulation on ALM activity (related to Figure 3) (A) Schematic of experiments. The anterolateral motor cortex (ALM) was recorded in L7-ChR2 mice preforming the task during photoactivation of Purkinje cells (PCs) in lateral crus 1 either unilaterally (only left crus 1) or bilaterally. (B) From top to bottom, average (line) and standard deviation (shaded area) of running speed, lick rate, and type 1-3 ALM neuronal activity during control (grey), unilateral crus 1 (blue), or bilateral crus 1 photostimulation (red) aligned on photostimulation onset (vertical dotted line). The laser activation profile is illustrated above the running speed traces. A ramping offset was introduced to limit the rebound of neuronal activity. (C) Average response profiles of firing rate from type 1-2 ALM neurons with significant reduction in activity compared to control trials, during the first 30 milliseconds following PC photostimulation onset (vertical dotted line) in unilateral (blue) or bilateral crus 1 photostimulation trials (red). Shaded areas represent SEM. Color-coded stars indicate first bin where firing rate significantly deviates from baseline (8 ms for unilateral crus 1 photostimulation, 14 ms for bilateral crus 1 photostimulation; see Methods).

**Figure S11.**
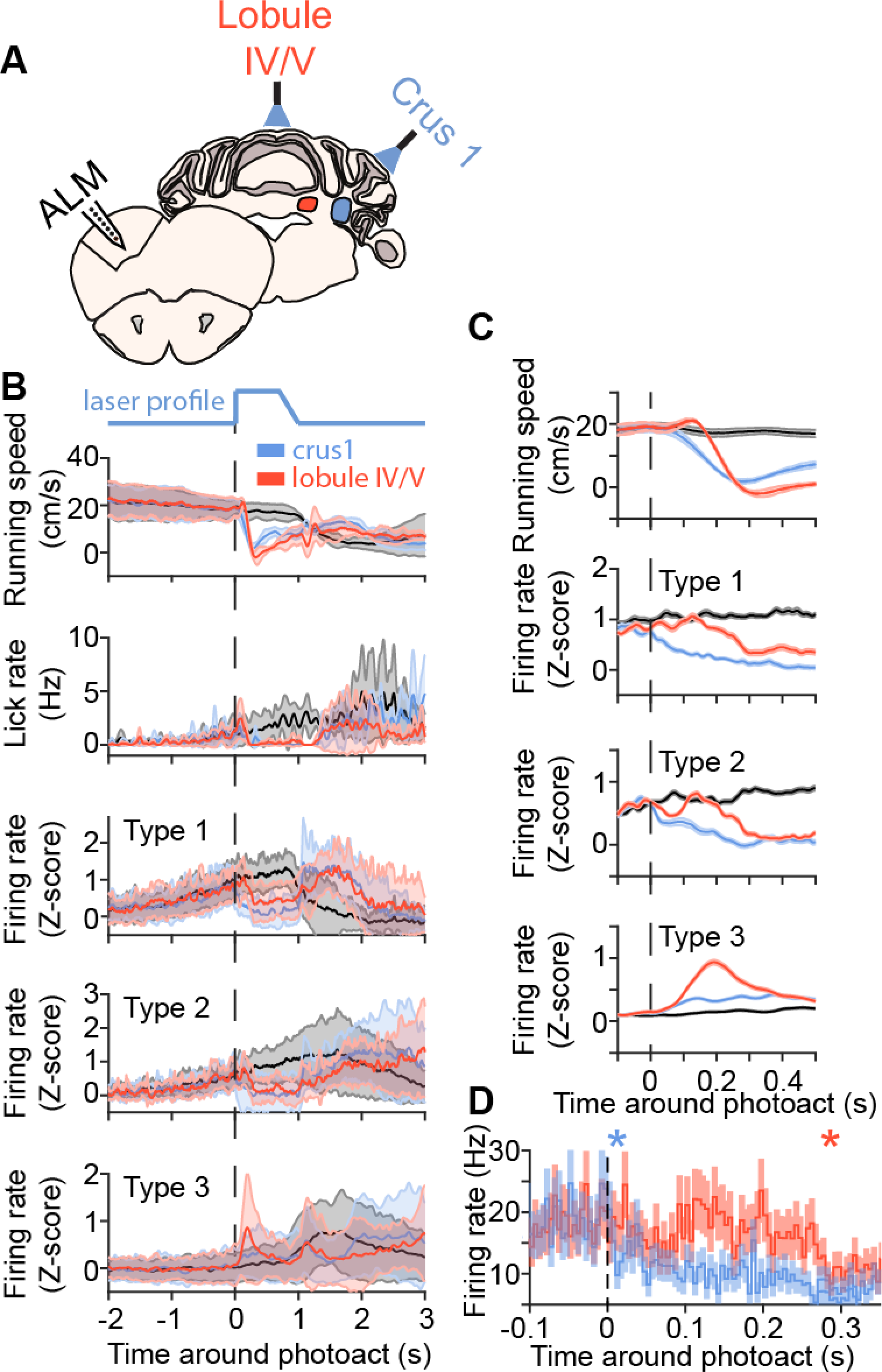
Comparison of the effect of lateral crus 1 and lobule IV/V photostimulation on ALM activity (related to Figure 3) (A) Schematic of experiments. The anterolateral motor cortex (ALM) was recorded in L7-ChR2 mice preforming the task during photoactivation of Purkinje cells (PCs) in lateral crus 1 or lobule IV/V. Lateral crus 1 PCs inhibit the dentate nucleus (blue region) and lobule IV/V PCs inhibit the fastigial nucleus (red region). (B) From top to bottom, average (line) and standard deviation (shaded area) of running speed, lick rate, and type 1-3 ALM neuronal activity during control (grey), lateral crus 1 (blue), or lobule IV/V photostimulation (red) aligned on photostimulation onset (vertical dotted line). The laser activation profile is illustrated above the running speed traces. A ramping offset was introduced to limit the rebound of neuronal activity. (C) From top to bottom, average (line) and SEM (shaded area) of running speed, and type 1-3 ALM neuronal activity during the first 500 milliseconds following photostimulation onset. (D) Average response profiles of firing rate from type 1-2 ALM neurons with significant reduction in activity compared to control trials, during the first 350 milliseconds following PC photostimulation onset (vertical dotted line) in lateral crus 1 (blue) or lobule IV/V (red). Shaded areas represent SEM. Color-coded stars indicate first bin where firing rate significantly deviates from baseline (10 ms for lateral crus 1 photostimulation, 280 ms for lobule IV/V photostimulation; see methods).

**Figure S12.**
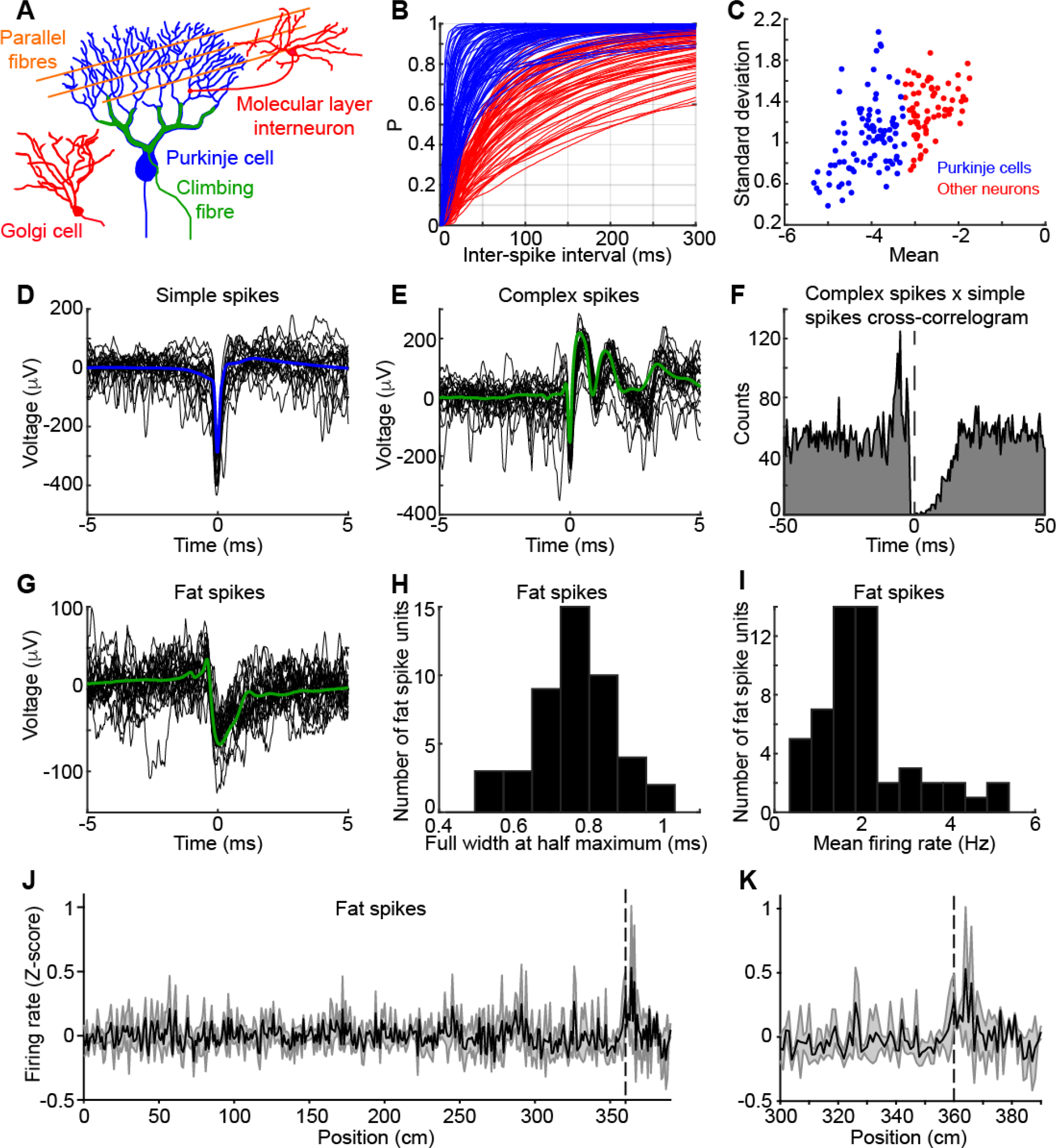
Single unit classification in the cerebellar cortex (related to Figures 4 and 5) (A) Schematic drawing of cerebellar cortex elements which activity can appear in our extracellular recordings. Purkinje neurons (PCs, blue) are inhibited by molecular layer interneurons (red) and have two sources of inputs: parallel fibres (orange) that give rise to simple spikes and climbing fibre (green) that give rise to complex spikes also recorded as fat spikes. Golgi cells are inhibitory interneurons found in the granule cell layer. (B) Cumulative distributions of inter-spike intervals (ISI) from units classified as PCs (blue) or other neurons (red). Units shown in this figure where recorded in crus 1 and 2. (C) ISI distributions were fitted with a log-normal function. K-means clustering with 2 clusters was performed on the mean and standard deviation of the ISI fits to separate putative PCs (blue dots) from other neurons. (D) Twenty successive (black) and average (blue) simple spike (SS) waveforms from a putative PC (high-pass filtered at 100 Hz). (E) Same as (D) for complex spikes (CSs) from the same PC. (F) Cross-correlogram (0.5 ms binning) between all SSs (examples in D) and CSs (examples in E). Note the characteristic pause in SS firing following the occurrence of a CS and lasting a few tens of milliseconds. All CSs belong to neurons classified as PCs in (C). (G) Same as (D) for a unit classified as a fat spike (FS; see Methods). (H) Distribution of FSs full width at half maximum. FSs duration is significantly longer than neuronal spikes and are likely to result from the large depolarisation of PCs dendritic trees induced by climbing fibre discharges. (I) Distribution of FSs mean firing rates. Note that those low firing rates are characteristic of climbing fibre activity. (J) Average firing rate (Z-score) +/− SD of all FSs spikes significantly modulated after reward delivery (*n* = 11/26; see Methods). FSs most regularly occur slightly further than the position of reward delivery (360 cm vertical dotted line). (K) same as (J) focusing on the rewarded section of the corridor.

**Figure S13.**
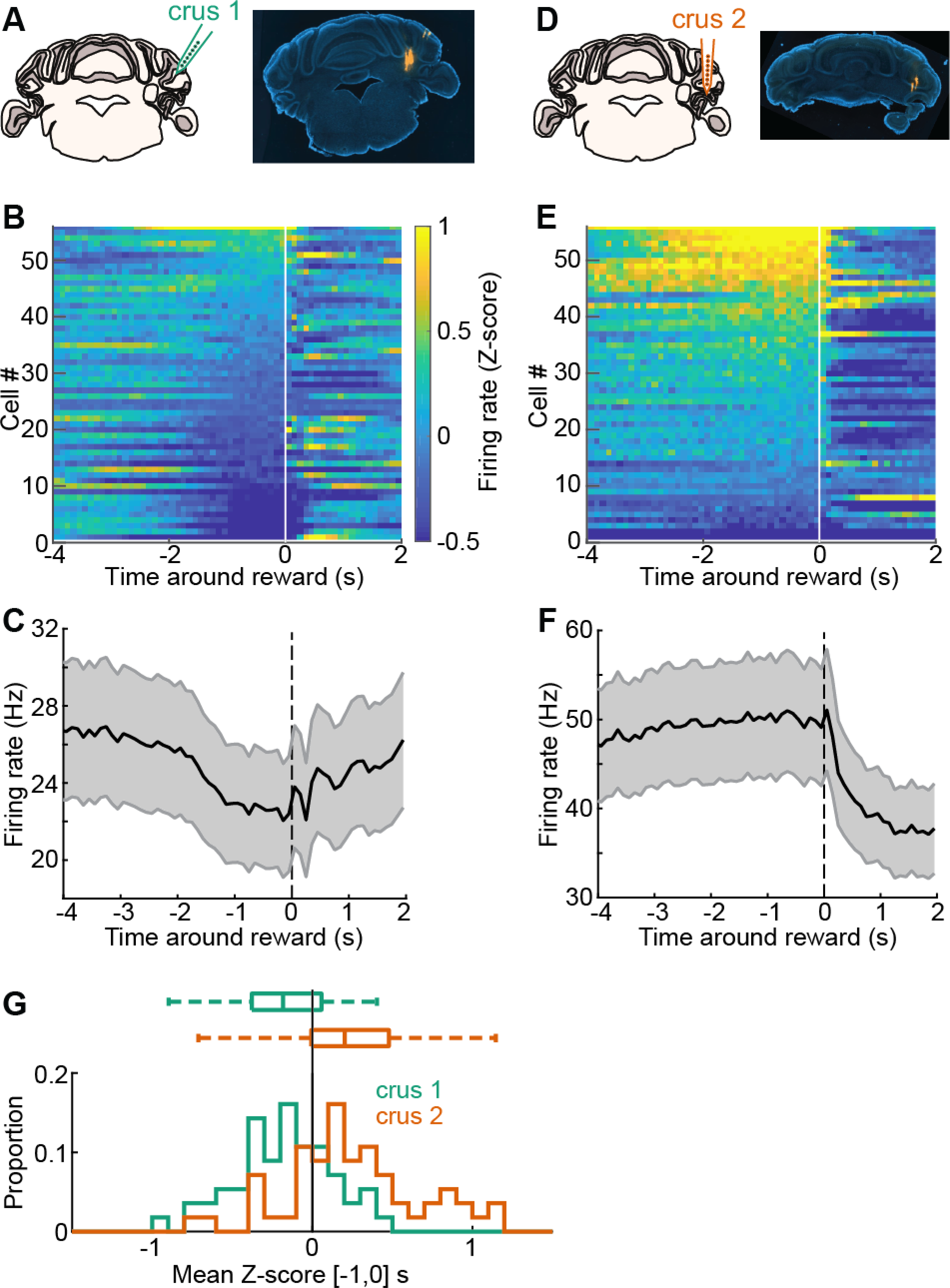
Purkinje cells in crus 2 show little sign of decrease in activity before reward unlike in crus 1 (related to Figure 4) (A) Left: Schematic of recording location in lateral crus 1. Right: Coronal brain slice from a lateral crus 1 recording (DiI tracks). Not that the DiI tracks in the DCN are from simultaneous DN recording. (B) Average response profiles for all crus 1 Purkinje cells (PCs) sorted by their mean Z-score value in the last second before reward. White vertical line indicates reward time. (C) Average PC population firing rate (black line) and SD (grey shaded area) showing decrease of activity starting around two seconds before reward (vertical dotted line). (D) Same as (A) for recordings in crus 2. (E) Same as in (B) for crus 2 PCs. (F) Same as in (C) for crus 2 PCs. (G) Distribution (bottom) and bar plot (top) of mean Z-score value in the last second before reward for crus 1 (green) and crus 2 PCs (orange; *p* < 0.0001, Wilcoxon Rank-Sum test).

**Figure S14.**
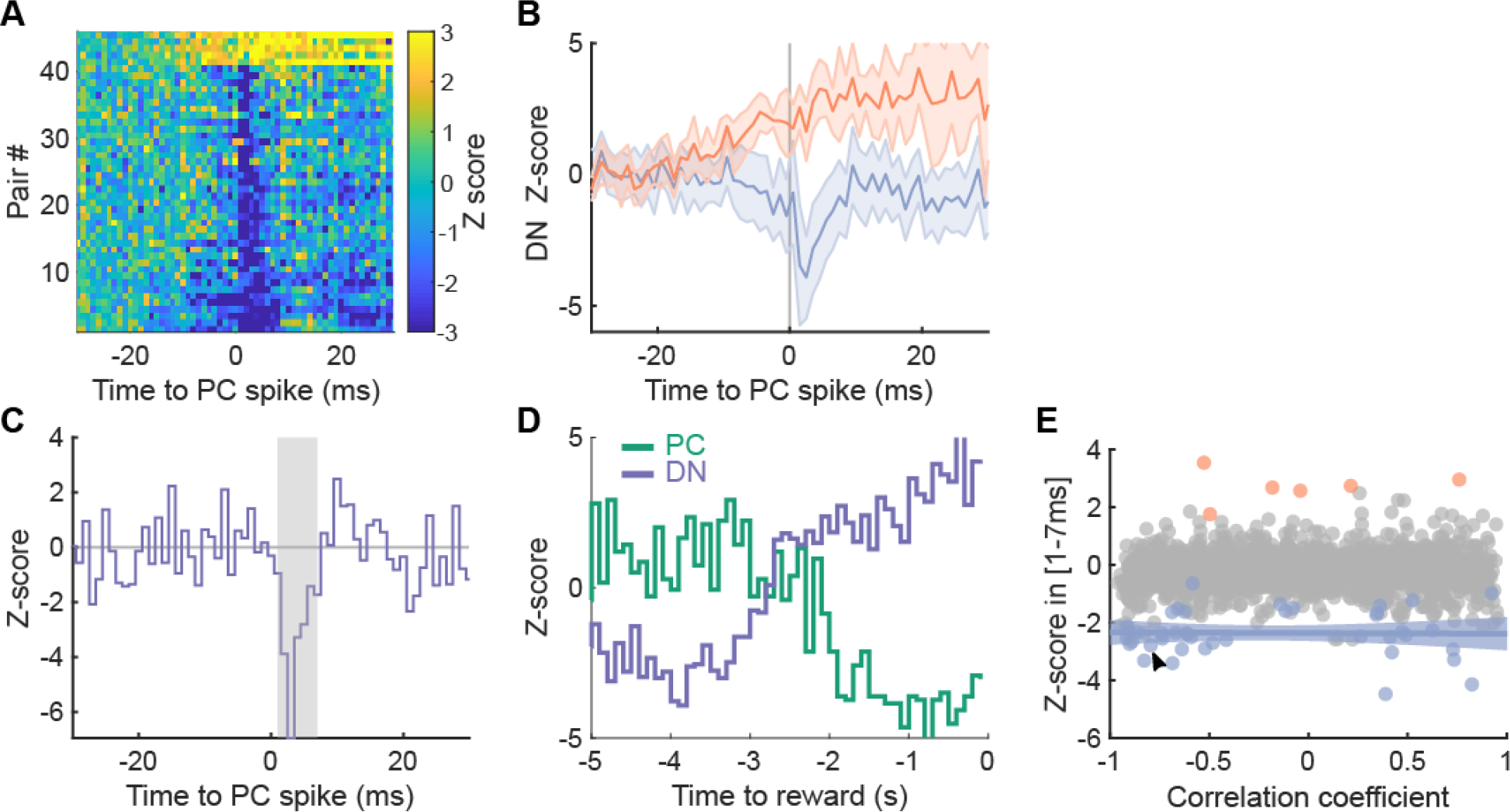
Functional connectivity between PCs and DN neurons (related to Figure 4) (A) Color-coded cross-correlograms (Z-score) between all modulated PC-DN neuron pairs (*n* = 46; see Methods). (B) Average peristimulus histogram +/− SD for excited (orange, *n* = 6) and inhibited pairs (blue, *n* = 40). (C) Cross-correlogram for an example PC-DN neuron pair. Grey shaded area indicates the time window for measuring the strength of connections (1 to 7ms after PC spike). (D) Z-score firing rate aligned to reward time (t = 0s) for the PC (green) and DN neuron (purple) of the pair shown in (C). (E) Strength of inhibition (measured as in (C)) versus correlation coefficient between the DN and PC peristimulus histograms in the 5 seconds before rewards (as in (D)). Blue dots are from inhibited pairs, orange dots are excited pairs, grey dots non-modulated pairs (see Methods). Blue line and shaded area, linear fit of inhibited pairs with 95% confidence interval. Arrowhead indicates example pair shown in (C) and (D).

**Figure S15.**
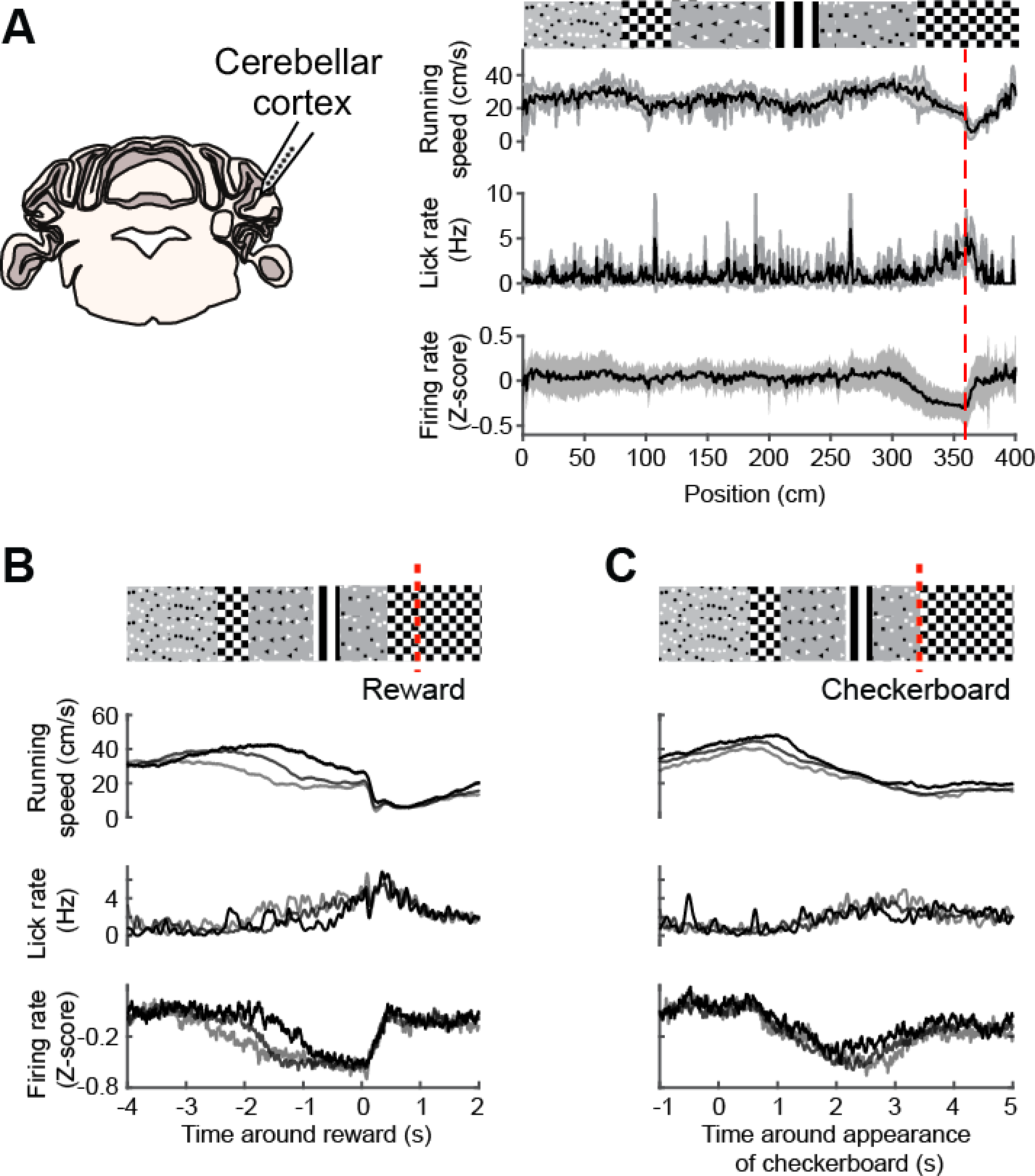
Lateral crus 1 PC activity mirrors preparatory activity in the reward context (related to Figures 4 and 5) (A) Left: schematic of recording location in lateral crus 1. Right: from top to bottom, average running speed, lick rate and activity of PCs exhibiting significant pre-reward reduction of firing rate (*n* = 25, 4 mice; see Methods) as a function of position inside the virtual corridor. Shaded areas represent SD. The visual patterns displayed on the corridor walls shown above the traces are aligned to the position at which they fully appear in the field of view of the mice (i.e. when they reach the back edge of the monitors). (B) From top to bottom, schematic of virtual corridor showing the location of reward delivery (red dotted line), mean running speed, lick rate, and PC firing rate binned according to the time at which mouse speed dipped under 20 cm/s in the 4 s before reward (1^st^ group : times below the 33^rd^ percentile of the distribution; 2^nd^ group: times between the 33^rd^ and 66^th^ percentile; 3^rd^ group: times above the 66^th^; *n* = 4 mice). (C) Same trial groups as in (D), aligned to the appearance of the rewarded checkerboard.

